# CryoEM and AI reveal a structure of SARS-CoV-2 Nsp2, a multifunctional protein involved in key host processes

**DOI:** 10.1101/2021.05.10.443524

**Authors:** Meghna Gupta, Caleigh M. Azumaya, Michelle Moritz, Sergei Pourmal, Amy Diallo, Gregory E. Merz, Gwendolyn Jang, Mehdi Bouhaddou, Andrea Fossati, Axel F. Brilot, Devan Diwanji, Evelyn Hernandez, Nadia Herrera, Huong T. Kratochvil, Victor L. Lam, Fei Li, Yang Li, Henry C. Nguyen, Carlos Nowotny, Tristan W. Owens, Jessica K. Peters, Alexandrea N. Rizo, Ursula Schulze-Gahmen, Amber M. Smith, Iris D. Young, Zanlin Yu, Daniel Asarnow, Christian Billesbølle, Melody G. Campbell, Jen Chen, Kuei-Ho Chen, Un Seng Chio, Miles Sasha Dickinson, Loan Doan, Mingliang Jin, Kate Kim, Junrui Li, Yen-Li Li, Edmond Linossi, Yanxin Liu, Megan Lo, Jocelyne Lopez, Kyle E. Lopez, Adamo Mancino, Frank R. Moss, Michael D. Paul, Komal Ishwar Pawar, Adrian Pelin, Thomas H. Pospiech, Cristina Puchades, Soumya Govinda Remesh, Maliheh Safari, Kaitlin Schaefer, Ming Sun, Mariano C Tabios, Aye C. Thwin, Erron W. Titus, Raphael Trenker, Eric Tse, Tsz Kin Martin Tsui, Feng Wang, Kaihua Zhang, Yang Zhang, Jianhua Zhao, Fengbo Zhou, Yuan Zhou, Lorena Zuliani-Alvarez, QCRG Structural Biology Consortium, David A Agard, Yifan Cheng, James S Fraser, Natalia Jura, Tanja Kortemme, Aashish Manglik, Daniel R. Southworth, Robert M Stroud, Danielle L Swaney, Nevan J Krogan, Adam Frost, Oren S Rosenberg, Kliment A Verba

## Abstract

The SARS-CoV-2 protein Nsp2 has been implicated in a wide range of viral processes, but its exact functions, and the structural basis of those functions, remain unknown. Here, we report an atomic model for full-length Nsp2 obtained by combining cryo-electron microscopy with deep learning-based structure prediction from AlphaFold2. The resulting structure reveals a highly-conserved zinc ion-binding site, suggesting a role for Nsp2 in RNA binding. Mapping emerging mutations from variants of SARS-CoV-2 on the resulting structure shows potential host-Nsp2 interaction regions. Using structural analysis together with affinity tagged purification mass spectrometry experiments, we identify Nsp2 mutants that are unable to interact with the actin-nucleation-promoting WASH protein complex or with GIGYF2, an inhibitor of translation initiation and modulator of ribosome-associated quality control. Our work suggests a potential role of Nsp2 in linking viral transcription within the viral replication-transcription complexes (RTC) to the translation initiation of the viral message. Collectively, the structure reported here, combined with mutant interaction mapping, provides a foundation for functional studies of this evolutionary conserved coronavirus protein and may assist future drug design.

## Introduction

Upon entry into human cells SARS-CoV-2, the causative agent of COVID-19, produces two large polyproteins, pp1a and pp1ab. These polyproteins are further processed by two viral proteases into 16 individual non-structural proteins (nsp1-nsp16). These non-structural proteins fulfill a number of essential viral functions including RNA replication, replication proofreading, double-membrane vesicle formation, and others^1^. Many of these also interact with host factors to effectively subvert the host cell to meet the virus’s needs^2^. Such subversion, for example, includes suppressing host innate immune responses, host translation, nuclear import, and other effects^3–5^. Despite their central importance to viral pathogenesis, many non-structural proteins remain structurally and functionally uncharacterized. Furthermore, the interactions between virus and host proteins are even less understood, with only a handful of unique viral-host protein complex structures available. As viruses often hijack central nodes in host cell pathways, studying viral-host interactions in molecular detail can lead to a better understanding of the mechanisms of viral pathogenesis and of the fundamental host cell processes the virus targets. In addition, viral-host protein complexes are an attractive target for antiviral therapeutics as they are less likely to accrue resistance. In this report we focus on one of the least studied SARS-CoV-2 proteins, Nsp2.

Neither the functions nor the structure of Nsp2 are known. In SARS-CoV-1, Nsp2 deletion leads to a defect in viral replication but still yields viable viruses^6^. Interestingly, expression of Nsp2 from an alternate site in the genome does not rescue this defect, likely indicating the importance of correct timing for Nsp2 expression^7^. A number of studies mapping host-viral interactomes of SARS-CoV-1/2 and MERS have identified host proteins that interact with Nsp2^8–11^. These studies have implicated Nsp2 in processes ranging from translation repression to endosomal transport, ribosome biogenesis, and actin filament binding. Nsp2 in SARS-CoV-2 and other coronaviruses have been observed to localize to endosomes and replication-transcription complexes (RTC); but it’s currently unclear what role Nsp2 plays at these sites^10,12^. Although Nsp2 is present in SARS-CoV-1, CoV-2, MERS—and in closely related coronaviruses in bats, pangolins and other animals—there is a considerable sequence variation across different species (Sup Fig 1). This degree of variability may indicate rapid Nsp2 adaptation under host-specific selection pressures. Furthermore, genome sequencing of SARS-CoV-2 variants during the COVID19 pandemic reveals sites under positive selection in Nsp2, suggesting host-specific human adaptation following successful zoonotic transfer^13^. Importantly, a specific mutation in Nsp2, T85I, is observed in clade 20C (as per Nexstrain nomenclature) including both variants of interest (B.1.526) and variants of concern (B.1.427/B.1.429 recently identified in California)^14,15^. Genetic knockdown/knockout studies have shown that a number of Nsp2 host interactors negatively affect SARS-CoV-2 replication, further corroborating the functional importance of Nsp2 ^10,16^.

Structural models of Nsp2 and Nsp2-host protein complexes will allow spatial mapping of the growing list of mutations and potentially shine light on their significance for Nsp2 functions. By delineating which Nsp2 surfaces determine specific Nsp2-host protein interactions, researchers will be able to generate specific Nsp2 point mutants that are defective in forming these protein complexes—helping tease apart the biological role of a particular Nsp2-host interaction. However, there are neither structures available for Nsp2 from any of the beta coronaviruses family members nor are there any similar structures in the PDB based on the sequence homology. Here we present a structure of SARS-CoV-2 Nsp2 derived from a combination of cryo-EM experimental data and AlphaFold2 prediction. Utilizing the natural sequence variation in SARS-CoV-2 together with our structure and mass spectrometry experiments, we identify two key Nsp2 surfaces that are required for specific host interactions.

## Results

### Combining a cryo-EM map with AlphaFold2 Nsp2 predictions yields a pseudo-atomic structure for full-length Nsp2

Full-length SARS-CoV-2 Nsp2 was recombinantly expressed and purified from *E*.*coli* cells. After purification involving multiple steps to remove contaminating nucleic acids, Nsp2 was obtained at high purity, plunge frozen, and imaged by cryo-electron microscopy. Processing in cryoSPARC2^17,18^ followed by RELION3^19^ yielded a reconstruction of Nsp2 at 3.8 Å global resolution ranging from 3Å in the best resolved regions to 6Å in the most flexible regions (Fig 1A and Sup Fig 2). The initial model was automatically built into the well-resolved region using the DeepTracer^20,21^ online server and then further corrected and refined via manual manipulation in Coot^22^ and ISOLDE^23^, followed by final refinements in Phenix^24^ and Rosetta^25^ (see methods). Having built an initial model, we then identified a number of putative zinc binding sites and repeated the sample purification and cryo-EM imaging in the presence of zinc. This yielded an improved 3.2 Å cryo-EM map, which revealed additional details and enabled improved modeling of residues 5-505 of the SARS-CoV-2 Nsp2 (Fig 1B). To our surprise, under these Zn-included conditions the density for the C-terminal 130 amino acids was completely missing. In the cryo-EM map without zinc, although the density for the flexible C-terminal domain was present, it was resolved at between 5-6 Å resolution. The closest homologous structure showed less than 10% sequence identity (PDB:3LD1)^26^, and as the domain was predicted to be high in the beta sheet fold this posed a significant challenge for *de novo* modeling based on low-resolution cryo-EM density alone.

**Figure 1.**
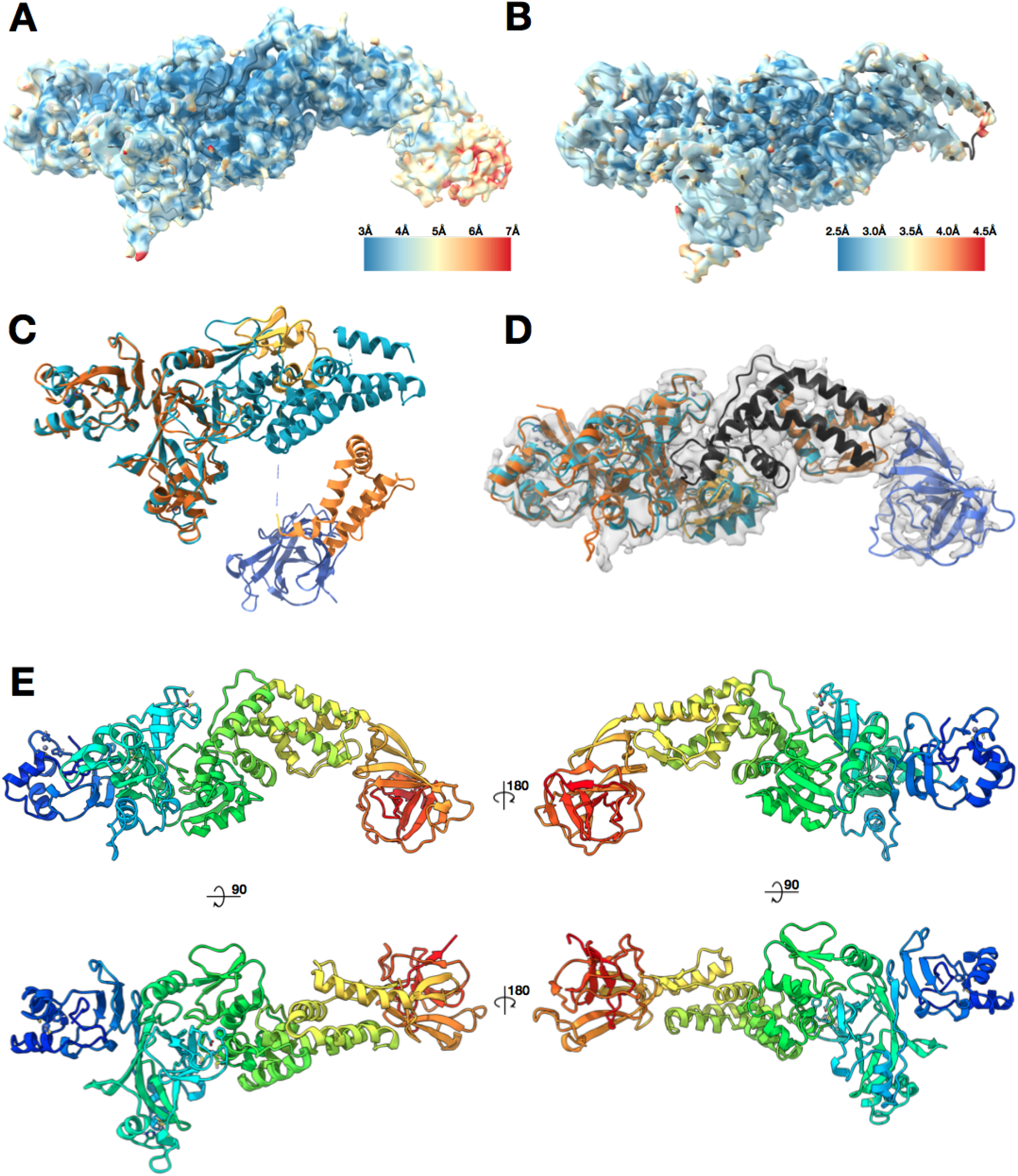
Nsp2 cryo-EM density and model overview. **(A)** 3.8 Å cryo-EM map of Nsp2 colored by local resolution showing the extra density at the C-terminus **(B)** 3.2 Å cryo-EM map of Nsp2 colored by local resolution with the resulting model in ribbon **(C)** Most up to date AlphaFold2 Nsp2 model (multicolored) was aligned to the experimentally built Nsp2 model shown in cyan ribbon. The missing 93 amino acids from the latest AlphaFold2 prediction are indicated by a dashed line. **(D)** AlphaFold2 Nsp2 predicted model (same as **C**) was broken into 4 regions and then individually aligned to the experimentally built model (domains segmented from the AlphaFold2 prediction are in shades of orange, experimental model is in cyan, in black is the region missing from the AlphaFold2 prediction but built into the experimental model, in blue is the C-terminal domain as predicted by AlphaFold2 and fit into the 3.8 Å cryo-EM map) (E) The resulting full length Nsp2 structure depicted as ribbon and colored as rainbow, blue for N terminus to red for C-terminus.

The recent utilization of deep learning for protein structure prediction based on amino acid sequence has led to a new level of success, as demonstrated by CASP14^27^. Specifically, the AlphaFold2 team was able to predict protein structures with unprecedented accuracy, producing results sometimes indistinguishable from the experimentally derived structures. AlphaFold2 and other teams in the CASP14 also ran predictions on SARS-CoV-2 proteins, including Nsp2. Out of all the available predicted models for Nsp2, only one model has an RMSD of less than 20 Å to our experimental model: C1901TS156_4 from the AlphaFold2 team. The other 5 models from AlphaFold2 were also close to 20Å RMSD, so we aligned all the available Nsp2 models from the AlphaFold2 team (5 from the CASP14 and one updated model available on their website^28^) to our structure (Sup Fig 3). This comparison made it clear that, globally, the predictions were quite different from the experimentally derived structure. In addition, the most updated model was missing a prediction for 93 amino acids of the protein (Fig 1C).

However, when analyzed in isolation, the individual motifs and domains of the proteins are remarkably close to the experimentally derived structure. This observation prompted us to break the model down into 4 subregions and align them to the experimentally derived structure independently. This yielded high local similarity per domain (average RMSD values of less than 2 Å, Fig 1D). The prediction for the missing C-terminal 130 amino acids in isolation fit well within the lower resolution density for that domain in the cryoEM map without zinc. We therefore combined the AlphaFold2 domain prediction for the C-terminal 130 amino acids with our experimentally built cryo-EM model to yield an experimentally valid and complete structure of full-length Nsp2 (Fig 1E, Sup Table 1).

### Nsp2 shows low global conservation among beta-coronaviruses, but possesses a highly conserved Zn binding motif

To better understand which regions of Nsp2 are functionally important, we performed a sequence alignment of Nsp2 across beta-coronaviruses from different species. Nsp2 shows low conservation, with the N-terminal half of the protein being marginally more conserved (Fig 2A and Sup Fig 1). Overall, SARS-CoV-2 Nsp2 is 68% identical to SARS-CoV-1 Nsp2 and only 20% identical to MERS virus Nsp2. Strikingly, the most conserved residues are a cysteine quad coordinating a Zn^2+^ ion in a Zn ribbon like motif, with three of the four cysteines being invariant across all the virus sequences. Performing a structural similarity search with this motif from Nsp2 indicates that it is similar to zinc ribbons^29^ in a number of RNA binding proteins in RNA polymerases and ribosomes (Fig 2A insert, average RMSD for the region of 1.7 Å). In some of these proteins these motifs explicitly have been implicated in RNA binding and in one structure (PDB:1JJ2, chain 2), the zinc ribbon on the ribosomal protein L44E is directly interacting with the ribosomal RNA. This motif is also similar to the tudor domains in the histone tail binding protein JMJD2A (RMSD of 1.7 Å). Although the fold is similar, the tudor domain is missing a Zn^2+^ ion in the JMJD2A structure (PDB:2QQS). Previous studies have associated Nsp2 with the viral RTCs^12,30^ and during the purification from bacteria we observed strong, apparently non-specific binding to *E*.*coli* nucleic acids that required chromatographic separation. One possibility, therefore, is that this motif is important for Nsp2 interactions with nucleic acids.

**Figure 2.**
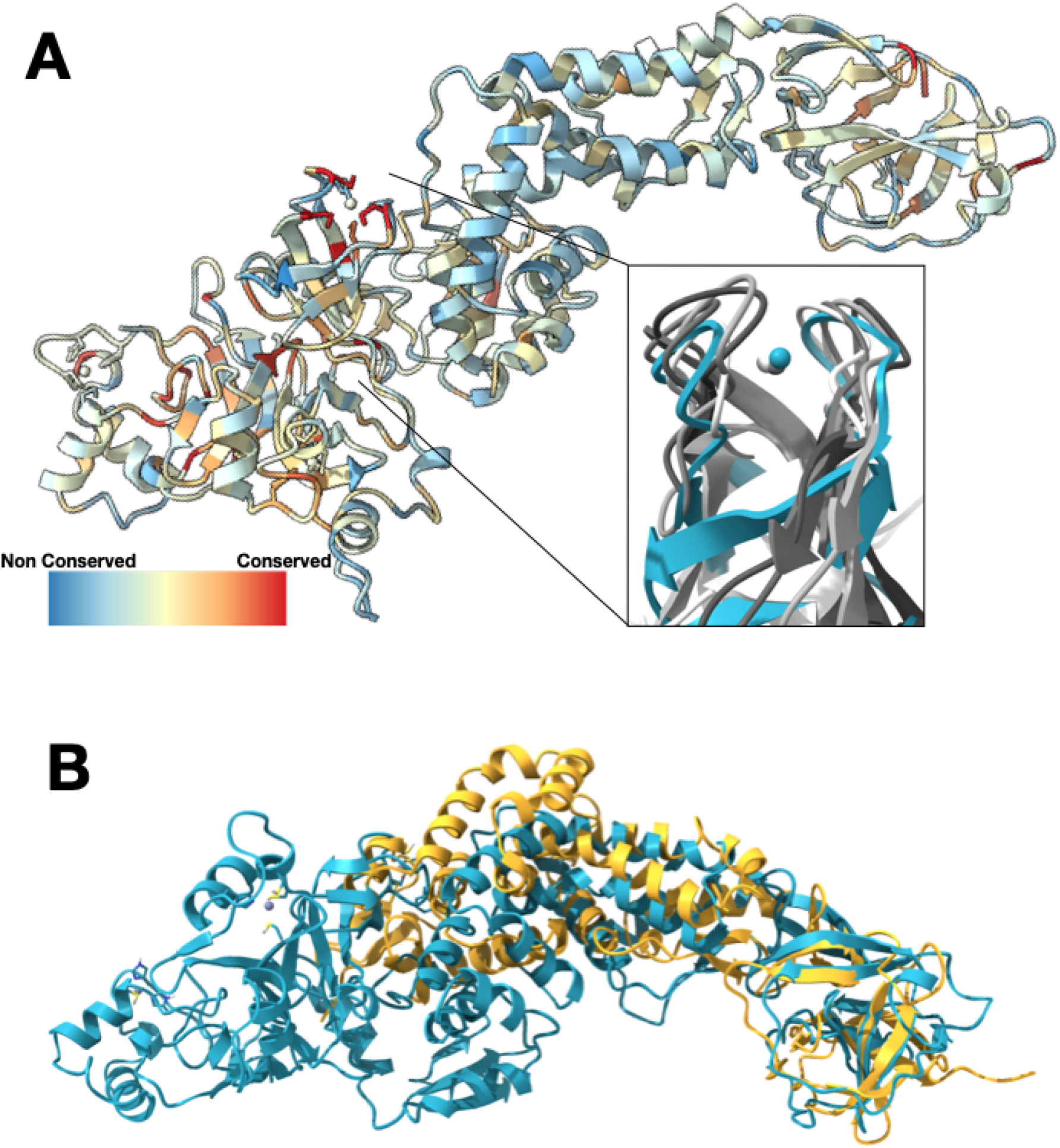
Nsp2 has a conserved zinc binding motif but otherwise shows low conservation. **(A)** Nsp2 structure depicted as ribbon and colored by conservation (see methods for details). The four cysteines show the highest conservation and are indicated in red. The magnified insert shows the zinc ribbon motif of Nsp2 in cyan aligned to zinc ribbon motifs from structurally similar structures in the PDB in shades of gray (PDBs: 1JJ2, 5XON, 1QUP, 4C2M). **(B)** Structure of IBV Nsp2 (PDB:3LD1, yellow ribbon) aligns well to the C-terminal region of SARS-CoV-2 Nsp2 (cyan) even though it has less than 10% sequence identity.

In addition to the proteins containing zinc ribbons and tudor motifs, a search of the PDB for structurally similar proteins returned only one additional structure, the structure of Nsp2 from Avian Infectious Bronchitis virus (PDB:3LD1). Although the sequence identity is below 10% for these proteins, the beta sheet C-terminal domain aligns well with our model. No other structures came up in our structural similarity search with either the FATCAT^31^ or DALI ^32^servers.

### Subsets of acquired mutations in Nsp2 group into surface patches

Examining the mutations that occur in SARS-CoV-2 Nsp2 during the COVID19 pandemic, over 50 sites have been identified as being under positive selection (at the time of writing, based on the dn/ds>1 metric^33,13^). Most of these mutations occur at low frequency. Two mutations however, T85I and I120F, are present at frequencies of roughly 13% and 5% respectively. The T85I mutation maps to a surface residue on our structure (Fig 3). The side chain of T85 is surface exposed, therefore replacing it with a hydrophobic isoleucine should not be favorable. However, if this region of Nsp2 is involved in protein-protein interactions such a substitution might be a gain-of-function change, stabilizing a hydrophobic binding interface. The second residue that is mutated, I120F, is not surface exposed and instead packs in a hydrophobic core that anchors a small helix. This small helix is attached to a highly charged loop on the surface of the protein and its dynamics may be functionally important. A phenylalanine mutation may further stabilize this helix anchor point by participating in pi-stacking interactions with neighboring aromatic residues (Fig 3 inset).

**Figure 3.**
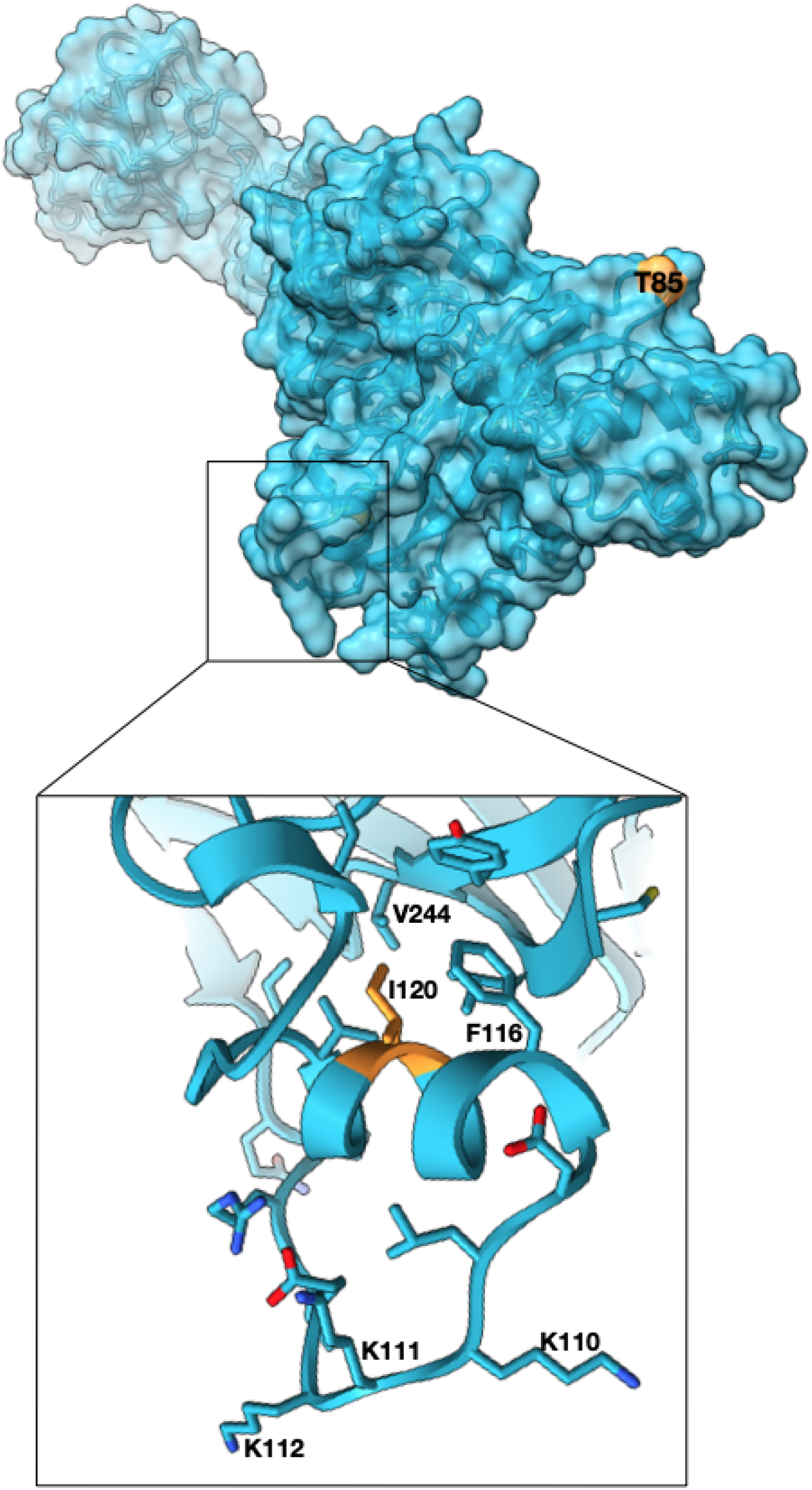
High frequency mutations in Nsp2 may provide host specific advantages. T85I mutation which is present in 13% of all the SARS-CoV-2 sequences is at the surface and may mediate host specific protein-protein interactions (Nsp2 surface in cyan, T85 in orange). Another mutated site, I120 points into a hydrophobic core stabilizing a small helix which is attached to a highly positively charged surface loop. Phe substitution at the site may further stabilize the helix. (zoomed panel, I120 in orange).

The structure allowed us to map the spatial relationships of conserved residues in Nsp2 among SARS-CoV-2 strains, revealing unexpected regions of conservation and selection. To identify rapidly evolving regions of the protein, we mapped all the positively-selected mutations to the protein surface (Fig 4). This analysis revealed charged surfaces which are devoid of mutations, potentially indicating surfaces important for conserved interactions (Fig 4). There are also two residue clusters where mutations found in strain variants are proximal to one another and alter the characteristics of the protein’s surface in similar ways. Cluster 1 is near the N-terminus consisting of three arginine residues (R27C, R52C, R4C) that mutate individually to cysteines, reducing the exposed positive surface charge in that region and introducing a sulfhydryl. Cluster 2 consists of six proximal residues which mutate individually to more hydrophobic residues (G262V, G265V, G285V, A411V, T371I) (Fig 4) in the variant strains. In both clusters, only individual single residue mutations are picked up in the viral population but the fact that they form physical clusters and have similar biochemical consequences might indicate adaptation.

**Figure 4.**
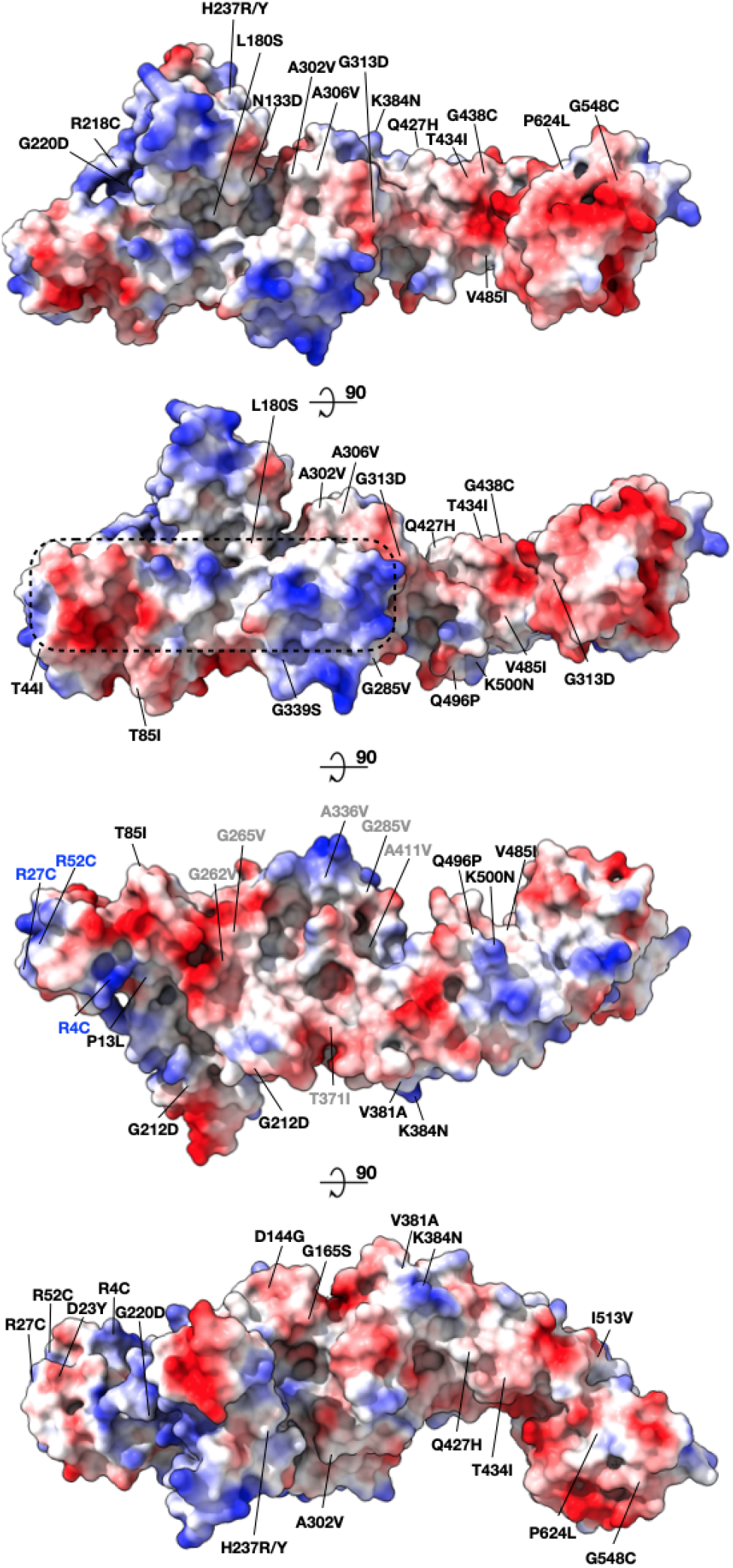
Mapping surface mutations in SARS-CoV-2 Nsp2 shows both potentially constrained surfaces and rapidly changing regions. All the positively selected mutations on Nsp2 mapped to the protein surface are colored by the surface charge. The region that is less susceptible to mutations is inscribed in a dashed rectangle. Residues that became less charged at the N-terminus are marked in blue (cluster 1). Residues that became more hydrophobic are marked in gray (cluster 2).

### Affinity purification mass spectrometry identifies Nsp2 surfaces mediating specific host interactions

To investigate whether common strain variants as well as disruptions of conserved patches would affect Nsp2’s ability to interact with host proteins we generated appropriate Nsp2 mutants and assessed changes in virus-human protein-protein interactions by affinity purification mass spectrometry (AP-MS) in HEK293T cells^2^. Three Nsp2 mutants were based on the natural Nsp2 variations: T85I, D23Y/R27C (cluster 1), G262V/G265V (cluster 2) and two were designed to disrupt the conserved/charged surface patches: K330D/K337D and E63K/E66K. All mutants expressed at similar levels in HEK293T cells (Sup Fig 4). We began by performing a global and unbiased quantification of virus-host protein-protein interactions using an automated affinity purification mass spectrometry workflow (see Methods). T85I and D23Y/R27C did not show significant changes in their interactomes, but three mutants did show significant changes in host interactions: G262V/G265V mutations abrogated Nsp2 interactions with the complex comprising GIGYF2/EIF4E2/RNF598^34^, E63K/E66K reduced interactions between Nsp2 and WASHC4/WASHC5 complex and also FKBP15, and K330D/K337D had a severely reduced interaction with NADPH cytochrome P450 reductase (POR gene) (Figure 5A,B,D, Table S2). Interestingly, Nsp2 E63K/E66K also gained a large number of new interactors which are predominantly involved in ribosomal RNA metabolic processes (Sup Fig 5). To increase the sensitivity and robustness of our quantitation, we performed Parallel Reaction Monitoring (PRM) on the subset of significantly changed interactors from the Data Dependent Analysis (DDA). Overall the PRM analysis recapitulated the findings of the DDA (Figure 5C).

**Figure 5.**
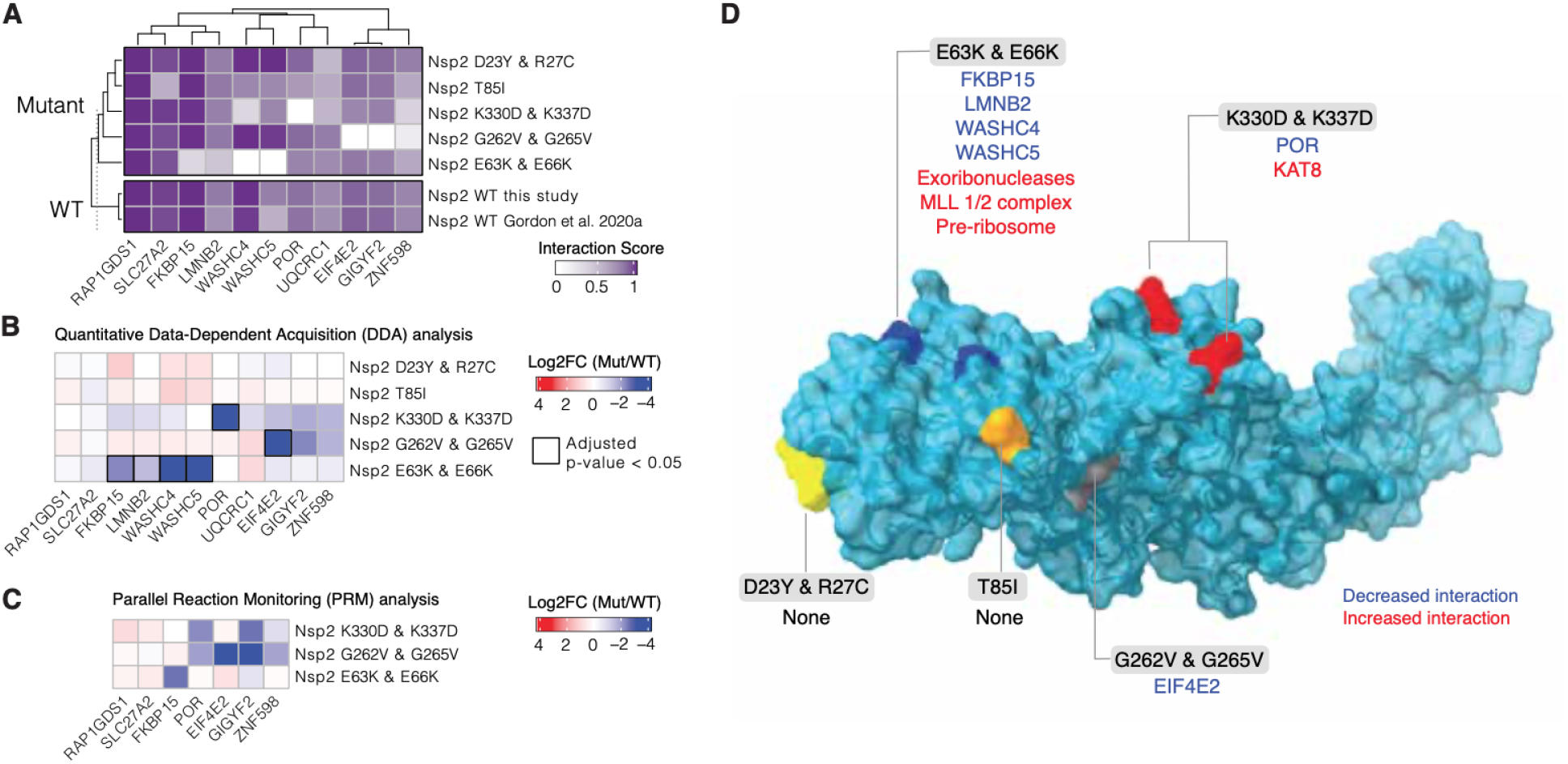
Nsp2 possesses multiple interaction surfaces for host proteins. **(A)** Interaction scores (average between MiST and Saint Scores) for human proteins (“preys”) deemed high-confidence interactions in at least one affinity purification (“bait”) mass spectrometry assay and detected to interact with both the wild-type Nsp2 in this study and in Gordon et al (2020a). Interaction scores range from zero to one, one being the most high-confidence. **(B)** Quantitative statistical analysis of data-dependent acquisition (DDA) mass spectrometry data using MSstats for interactions selected and depicted in **A**. Prey intensities were normalized by bait expression abundance. Log_2_ fold changes and BH-adjusted p-values were calculated by comparing each mutant to the wild-type from this study. Square black outlines depict adjusted p-values < 0.05. **(C)** Parallel reaction monitoring (PRM) analysis of select preys from B for mutants found to possess significantly-changed interactions (adjusted p-value < 0.05). **(D)** Nsp2 structure depicted as surface (light blue) with the mutations considered in this study depicted on the surface: E63K/E66K (dark blue), K330D/K337D (red), D23Y/R27C (yellow), T85I (orange), and G262V/G265V (grey). Lost interactions (adjusted p-value < 0.05) from data-dependent acquisition global proteomics analysis (DDA) from B depicted in blue and gained protein complexes depicted in red (see Sup Fig 4)

## Discussion

In this report, we were able to combine cryoEM with recent advances in de novo protein predictions to obtain a complete atomic model for SARS-CoV-2 Nsp2 protein. Although there was a recent report of using AlphaFold2 predicted protein structure of Orf8 to solve the phase problem in crystallographic studies, to our knowledge this is the first explicit use of AlphaFold2 predictions with restraints from an experimental cryoEM density for model building^35^. This exercise suggests that domain structure predictions from deep neural networks are increasingly likely to be locally accurate and, when combined with experimental restraints, sufficient for global structure prediction and integrative structural modelling. Electron cryo-microscopy and cryo-tomography will be important sources of such overall shape information, and readily obtainable, low-resolution measurements like negative stain electron microscopy, small-angle X-ray scattering, cross-linking mass spectrometry, or even biochemical experiments may provide sufficient constraints for accurate, global models to be determined in combination with predicted domain structures. It is possible that further improvements in the prediction algorithms will eliminate the need for experimental measurements entirely. However, atomic resolution structures of multi-component and multi-domain assemblies are still relatively uncommon, and this deficit of appropriate training data in the PDB may limit the accuracy of computational models for multi-domain assemblies and higher-order complexes. Put another way, the deficiency of data about protein-protein interfaces may mean that de-novo predictions of complex assemblies will remain underdetermined for some time. Future work will explore the use of restraints from 3D cryo-EM maps, 2D images, tomograms, and other data sources like SAXS for the potential functions utilized by neural nets.

Our Nsp2 structure together with analysis of natural and designed sequence variation in the Nsp2 of SARS-CoV-2 suggests a number of biological roles for Nsp2 and also regions of interest on the protein. We identify a highly conserved zinc ribbon motif which structurally is highly similar to zinc ribbons in RNA binding proteins. One possibility, therefore, is that this motif is important for Nsp2 interactions with nucleic acids. Interestingly our mass spectrometry studies on the E63K/E66K mutant, designed to introduce a charge reversal mutation in a conserved negatively charged surface patch (Figs 4,5 and Sup Fig 5), show that this mutant gains a large number of partners involved in ribosome biogenesis. Concurrently with this gain, this mutant loses interactions with the endosomal/actin machinery (FKBP15, WASHC proteins^36,37^). It is tempting to speculate that Nsp2 binds ribosomal RNA (via the identified zinc ribbon) but attachment to cytoskeletal elements at endosomes limits its interactions with rRNA. Upon disruption of the endosomal anchoring of Nsp2 by the above mutations, Nsp2 is free to interact with ribosomal RNAs—potentially even translocating to the nucleolus. Interestingly, previous proximity labeling studies on murine coronavirus Nsp2 have uncovered an exciting link between the viral polymerase within RTCs and the host cell’s translation machinery^12^. Since Nsp2 has affinity for ribosomal RNA and is localized to RTCs, one appealing model is that Nsp2 plays a role in locally enriching ribosomes next to viral messages for more efficient transcriptional-translational coupling in the cytosol. Another aspect of these specific interactions are potential roles that Nsp2 may play in hijacking the endosomal pathway to meet viral needs. WASH complexes have been shown to play key roles in exocytosis and endosome biogenesis, including cargo sorting through local Arp2/3 complex activation^38^. SARS-CoV-2 enters cells through the endosomal pathway and it may be functionally important for the virus to modulate endosome pathways for successful infection.

Mass spectrometry analysis of the second Nsp2 mutant, G262V/G265V implicates Nsp2 in modulating ribosome-associated quality control. This mutant is based on the natural variants of Nsp2 in the patch that is becoming more hydrophobic during the 2019-2020 COVID19 pandemic (Figure 4, cluster 2), although the natural variants all have a single mutation, either the G262V or G265V. This mutant is the only one that specifically loses interactions with GIGYF2, EIF4E2 and RNF598. These three proteins are known to form a complex and have been implicated in inhibiting translation initiation when ribosomes stall on defective or abnormal mRNA messages^34^. At the moment it is unclear what is the exact functional connection between SARS-CoV-2 infection and this recently discovered component of ribosome-associated quality control mediated by GIGYF2/EIF4E2. Given that SARS-CoV-2 has optimized codon usage for the human host, it is unlikely that there is increased ribosomal stalling on the viral message that requires inhibition of GIGYF2/EIF4E2 by Nsp2^39,40^. Furthermore, genetic studies have shown that GIGYF2 and EIF4E2 are necessary rather than inhibitory for viral replication ^16^. Perhaps the virus uses Nsp2 to inhibit translation initiation of *host* messages, freeing ribosomes and the rest of the translation machinery for virus production. Indeed, a prior study of the GIGYF2/EIF4E2/ZNF598 complex suggests that in addition to its role in RQC, this complex selectively suppresses transcripts involved in host inflammatory signaling, including IL-8^41^. It is worth noting that Nsp2 interactions with GIGYF2/EIF4E2/ZNF598 complex is conserved across SARS-CoV-1 and MERS indicating that this interaction might be of therapeutic importance for coronaviruses generally ^10^.

Our mass spectrometry experiments of the most prevalent Nsp2 mutation, T85I, did not identify any changes in host interactions of this mutant. This may be due to our experiments lacking the context of other viral proteins that would be present in a bona fide infection or potentially due to the wrong cellular context. Alternatively this may suggest that some mutations do not confer any fitness benefit and are simply present due to the C-U hypermutation observed in SARS-CoV-2, which is likely driven by host mediated APOBEC editing^42^. Interestingly, there is a recent report demonstrating that the SARS-CoV-2 Nsp2 T85I mutant shows a minor replication defect in Vero green monkey cells, but has no effect in human cells. This is consistent with the T85I mutation not conferring a strong selective advantage^43^. Globally, the second most prevalent SARS-CoV-2 amino acid substitution that is driven by the C-U hypermutation is a T to I change. Therefore the T85I mutation in the 20C clade of SARS-CoV-2 may be neutral in fitness, but stable due to host-mediated RNA editing.

Overall, analysis of the resulting Nsp2 structure revealed a rapidly evolving protein surface, with potential consequences for host-virus interactions. Leveraging the new structure with natural Nsp2 variations and mass spectrometry we were able to identify surfaces important for specific Nsp2 interactions. The pattern of disruption of interactions points to at least three specific areas of biology that Nsp2 is involved in: interactions with endosomes through cytoskeletal elements, interactions with modulators of translation, and also direct interactions with ribosomal RNA. The exact roles Nsp2 plays in these pathways will require further experimental characterization using the structure-based point mutants described here.

## Methods

### Nsp2 Expression

SARS-CoV-2 Nsp2, codon optimized for *Escherichia coli* expression, was cloned in a pET-29b(+) vector backbone with N-terminus 10XHis-tag and SUMO-tag (Ep156). For expression, the plasmid Ep156 was transformed in the LOBSTR *E. coli* strain and a single colony was inoculated in LB media with 50 µg/ml kanamycin overnight at 37°C. 1% overnight culture was inoculated in 1 L TB media with 50 µg/ml kanamycin and and grown at 37°C till O.D._600nm_ reached 0.8. The culture was transferred to 20°C and induced with 0.5 mM IPTG for 16 h. The cells were harvested and washed with PBS, flash-frozen and stored at −80°C.

### Nsp2 Purification

To a 6 L equivalent of cell pellet, 150 mL of lysis buffer (50 mM HEPES pH 7.5, 300 mM NaCl, 10% glycerol, 2 mM MgCl_2_) supplemented with 2 protease inhibitor tablets (Roche), 1 mM PMSF, and 30 μL benzonase nuclease (Millipore Sigma) was added. Cells were resuspended and dounce homogenized before either sonication (3 x 10 min cycles using a sonifier (Branson), at 50% duty cycle (1 sec on, 1 sec off), followed by ≥5 mins on ice, or high pressure homogenizer (3 times passage through the EmulsiFlex-C3 [Avestin] at ∼15,000 psi, 4°C). After centrifugation for 40 mins at 25,000xg, 4°C, clarified samples were transferred to a 50 mL conical tube and supplemented with a final 20 mM imidazole pH 7.5 before batch-binding with Ni-NTA superflow resin (Qiagen) for ∼1 hr at 4°C. The resins were collected in a gravity flow column, washed with 15 CV lysis buffer, followed by 2x 7.5 CV sulfate wash buffer (25 mM Tris pH 8.5, 300 mM NaCl, 10% glycerol, 100 mM Na_2_SO_4_), 2x 7.5 CV wash buffer (25 mM Tris pH 8.5, 300 mM NaCl, 10% glycerol, 30 mM imidazole), 2x 7.5 CV wash buffer supplemented with 2 mM ATP, 4 mM MgCl_2_, and eluted in 2x 2.5 CV elution buffer (25 mM Tris pH 8.0, 300 mM NaCl, 10% glycerol, 300 mM imidazole, 2 mM MgCl_2_). The elution was treated with benzonase and Ulp1 protease and dialyzed overnight at 4°C in dialysis buffer (25 mM Tris pH 8.5, 75 mM NaCl, 10% glycerol, 0.5 mM TCEP, 2 mM MgCl_2_). The tagless Nsp2 was further purified using a 5 mL HiTrap Heparin HP column (Cytiva) using a linear gradient of 7.5% Heparin buffer A (30 mM Tris pH 8, 1 mM DTT) to 50% Heparin buffer B (30 mM Tris pH 8, 1 M NaCl, 1 mM DTT). Peak fractions corresponding to Nsp2 were concentrated and further purified using a Superdex 200 increase 10/300 GL column (Cytiva) in SEC buffer (20 mM Tris pH 8.0, 250 mM NaCl, 0.5 mM TCEP) to yield a single peak. The peak fractions were pooled and concentrated and used for CryoEM. For the Zn containing sample, 10 µM ZnCl_2_ was kept in all the buffers of the purification protocol mentioned above.

### CryoEM grid freezing and data collection

Purified nsp2 was diluted to 6 µM for the apo sample and 5.7 µM for the Zn containing sample. 400 mesh 1.2/1.3R Au Quantifoil grids were glow discharged at 15 mA for 30 seconds. Vitrification was done using FEI Vitrobot Mark IV (ThermoFisher) set up at 4°C and 100% humidity. 3.5 µl sample was applied to the grids and the blotting was performed with a blot force of 0 for 4-6 s prior to plunge freezing into liquid ethane. For the apo sample, two datasets comprising of 804 and 1116 118-frame super-resolution movies each were collected with a 3×3 image shift at a magnification of 105,000x with physical pixel size of 0.834 Å/pix on a Titan Krios (ThermoFisher) equipped with a K3 camera and a Bioquantum energy filter (Gatan) set to a slit width of 20 eV. Collection dose rate was 8 e^-^/pixel/second for a total dose of 66 e^-^/Å^2^. Defocus range was 0.8 to 2.4 µm. Each collection was performed with semi-automated scripts in SerialEM. Nsp2 with Zn grids were prepared using a similar protocol. 1149 118-frame movies were collected for this sample at a 105,000x magnification with physical pixel size of 0.834 Å/pix on Titan Krios similar to without Zn sample. Collection dose rate was 8 e^-^/pixel/second for a total dose of 67 e^-^/Å^2^.

### Data Processing

#### Without Zn

Initial processing was done in Cryosparc (v2.15.0)^17 18^. The first dataset with 801 dose-weighted motion corrected micrographs^44^ was imported and Patch CTF(M) was performed. This dataset required thorough manual curation and led to selection of 388 micrographs for further processing. Blob-picker was used to pick 363145 particles and extraction was done with a box size of 288 px. 2D-classification was done into 150 classes and good looking classes were selected with total 91181 particles. A second dataset with 1116 dose-weighted micrographs was processed in a similar manner. After Patch CTF(M), 748 micrographs were curated based on CTF-fit resolution (<5 Å), ice-thickness, and carbon. Templates were created from the previous dataset and used for template-based particle picking to get 577518 particles. 240551 particles were selected after 2D-classification into 200 classes. These particles were merged with 91181 selected particles of the previous dataset after 2D-classification. Total 331732 particles were classified into 3 classes with heterogeneous refinement. Multiple rounds of heterogeneous refinements, non-uniform refinements and homogenous refinements resulted in a 3.45 Å 3D-reconstruction with 99076 particles. The particles from this final step of Cryosparc processing were imported into Relion^19^ (version 3.0.8) for further processing. 3D-classification without mask resulted in a 3.59 Å map. The core of the map was better resolved therefore skip-align classification was done on the core. The best class was subjected to 3D-refinement and post-processing leading to the final map at 3.49 Å. This class was selected and skip-align classification was done for the full map. The overall resolution of the full map with the selected 42579 particles was reported to be 3.76 Å which upon manual inspection was the best looking map even though nominally being at worse resolution than some previous reconstructions.

#### With Zn

Like the ‘without Zn’ dataset, for initial processing we used Cryosparc (v2.15.0). Patch CTF(M) was performed on imported 1149 dose-weighted micrographs. The micrographs were curated for CTF-fit resolution of better than 5 Å. Template-based particle picking was done (templates from the without Zn dataset) on the selected 1028 micrographs, resulting in 1515264 particles. A series of ab-initio classifications followed by heterogeneous refinements and non-uniform refinements on the best classes selected led to a map of 3.1 Å with 81817 particles. These particles were transferred to Relion (version 3.0.8) and a single-class skip-align classification was performed with a mask. A 3.15 Å map with 81817 particles was obtained after 3D-refinement and post-processing on the particles from skip-align classification.

### Refinement/Model building

The initial model for the core of ‘without Zn’ map was initially obtained by submitting the high resolution region of the map with the full Nsp2 sequence to the DeepTracer server^20 21^. This resulted in two chains which threaded the backbone fairly well but contained amino acid substitutions/deletions. Therefore a homology model was built for Nsp2 using the resulting model from DeepTrace server as a template, using SwissModel server. This model was then refined against the map in Phenix Real Space Refine^24^ and iteratively rebuilt with Rosetta (2020.08 release)^25^. Best scoring models were manually examined and corrected using COOT 0.9^22^. Rosetta was used for automatic iterative rebuilding the lower resolution regions of the map and loops (212-224, 475-490). Cys/His residues were manually identified for Zn coordination sites and Zn was placed in COOT. At this point the higher resolution cryo-EM map obtained in the presence of zinc was used for downstream steps. Map/model quality was examined and ramachandran outliers were fixed in ISOLDE 1.0^23^. Rosetta FastRelax was used in cartesian space followed by iterations of refinement in Phenix Real Space Refine. This fixed most of the geometry outliers but introduced a large number of clashes in the model. Rosetta FastRelax in torsion space was used on the model from Phenix Real Space Refine to resolve the clashes while preserving good model statistics for the final model. The b-factors were assigned using Rosetta B-factor fitting mover. Local resolution was determined by running ResMap program^45^. Directional FSC curves were determined by submitting the associated files to the 3DFSC server^46^.

### Obtaining full Nsp2 model incorporating AlphaFold2 prediction

Predicted Nsp2 models were downloaded either from the DeepMind website or CASP14 website and then were aligned to the experimental Nsp2 map using matchmaker tool in ChimeraX^47,48^. Based on visual examination, the most updated (at the time of writing) AlphaFold2 model (V3_4_8_2020) was split into 4 domains: 1-277, 278-344, 459-509 and 510-638. First three were individually aligned with matchmaker in ChimeraX to the experimental model to assess the similarity and report RMSDs in the main text. These regions were not used for downstream model building. The fourth region, 510-638 was rigid body fit into the 3.8 Å cryo-EM map (obtained without zinc). The experimental model was then stitched together with the rigid body fit domain for residues 510-638 from AlphaFold2. The whole model was energy minimized into the cryo-EM density filtered to 5 Å by running Rosetta FastRelax in torsion space.

### Sequence alignment and sequence conservation analysis

Nsp2 sequences were manually downloaded from UniProt and aligned in Jalview using MAFFT^49,50^. Conservation was mapped on the Nsp2 structure using a combination of Chimera and ChimeraX and the supplementary alignment figure was prepared with MView server^47,51,52^.

#### SARS-CoV-2 Nsp2 mammalian expression constructs

SARS-CoV-2 isolate 2019-nCoV/USA-WA1/2020 (accession MN985325), an early-lineage sequence downloaded on January 24, 2020, was the reference sequence for all viral expression constructs. Native nucleotide sequences encoding proteolytically mature Nsp2 were first codon optimized (https://www.idtdna.com/codonopt) for gene synthesis. The gBlock Gene Fragment (IDT) encoding Nsp2 and the C-terminal linker and 2x-Strep tag was inserted into pLVX-EF1alpha-IRES-Puro at the EcoRI and BamHI restriction sites by In-fusion cloning [PMID: 32353859]. SARS-CoV-2 Nsp2 mutants (D23Y/R27C, E63K/E66K, T85I, G262V/G265V, and K330D/K337D) were generated in a similar manner using unique restriction sites within the Nsp2 sequence to excise segments containing wild type residues. Sequences were subsequently replaced by In-fusion cloning with gBlocks (IDT) containing mutated residues. All mutations were confirmed by sequencing.

#### Cell culture

HEK293T cells were cultured in Dulbecco’s Modified Eagle’s Medium (Corning or Gibco, Life Technologies) supplemented with 10% Fetal Bovine Serum (Gibco, Life Technologies) and 1% Penicillin-Streptomycin (Corning) and maintained at 37°C in a humidified atmosphere of 5% CO_2_.

#### Transfection

For each affinity purification (wild-type and mutant nsp2 and controls, empty vector and EGFP), 7.5 million HEK293T cells were plated per 15-cm dish and allowed to recover for 20-24 hours prior to transfection. Up to 15 μg of individual Strep-tagged expression constructs (normalized to 15 μg with empty vector as needed) was complexed with PolyJet Transfection Reagent (SignaGen Laboratories) at a 1:3 μg:μl ratio of plasmid to transfection reagent based on manufacturer’s recommendations. After 40 hours, cells were dissociated at room temperature using 10 ml Dulbecco’s Phosphate Buffered Saline without calcium and magnesium (D-PBS) supplemented with 10 mM EDTA for at least 5 minutes and subsequently washed with 10 ml D-PBS. Each step was followed by centrifugation at 200 xg, 4°C for 5 minutes. Cell pellets were frozen on dry ice and stored at - 80°C. At least three biological replicates were independently prepared for affinity purification.

#### Affinity purification

Frozen cell pellets were thawed on ice for 15-20 minutes and suspended in 1 ml Lysis Buffer [IP Buffer (50 mM Tris-HCl, pH 7.4 at 4°C, 150 mM NaCl, 1 mM EDTA) supplemented with 0.5% Nonidet P 40 Substitute (NP40; Fluka Analytical) and cOmplete mini EDTA-free protease and PhosSTOP phosphatase inhibitor cocktails (Roche)]. Samples were then frozen on dry ice for 10-20 minutes and partially thawed in a 37°C water bath. Following two freeze-thaw cycles, samples were incubated on a tube rotator for 30 minutes at 4°C and centrifuged at 13,000 xg, 4°C for 15 minutes to pellet debris. After reserving 50 μl lysate, samples were arrayed into a 96-well Deepwell plate for affinity purification on the KingFisher Flex Purification System (Thermo Scientific) as follows: MagStrep “type3” beads (30 μl; IBA Lifesciences) were equilibrated twice with 1 ml Wash Buffer (IP Buffer supplemented with 0.05% NP40) and incubated with ∼ 0.95 ml lysate for 2 hours. Beads were washed three times with 1 ml Wash Buffer, once with 1 ml IP Buffer and then transferred to 75 μl Denaturation-Reduction Buffer [2 M urea, 50 mM Tris-HCl pH 8.0, 1 mM DTT) aliquoted into 96-well plates for on-bead digestion (below)]. The KingFisher Flex Purification System was placed in the cold room and allowed to equilibrate to 4°C overnight before use. All automated protocol steps were performed using the slow mix speed and the following mix times: 30 seconds for equilibration/wash steps, 2 hours for binding, and 1 minute for final bead release. Three 10 second bead collection times were used between all steps.

#### On-bead digestion

Bead-bound proteins were incubated in Denaturation-Reduction Buffer for 30 minutes, alkylated in the dark with 3 mM iodoacetamide for 45 minutes and quenched with 3 mM DTT for 10 minutes. Proteins were then trypsin digested as follows: initially for 4 hours with 1.5 μl trypsin (0.5 μg/μl; Promega) and then another 2 hours with 0.5 μl additional trypsin. To offset evaporation during trypsin digestion, 22.5 μl 50 mM Tris-HCl, pH 8.0 was added. All steps were performed with constant shaking at 1,100 rpm on a ThermoMixer C incubator set to 37°C (denaturation-reduction and trypsin digest) or room temperature (alkylation and quenching). Digested peptides were combined with 50 μl 50 mM Tris-HCl, pH 8.0 used to backwash beads and acidified with trifluoroacetic acid (0.5% final, pH < 2.0) Acidified peptides were desalted using a BioPureSPE Mini 96-Well Plate (20mg PROTO 300 C18; The Nest Group, Inc.) according to standard protocols and dried in a CentriVap Concentrator (Labconco) for at least two hours.

#### Mass spectrometry data acquisition and analysis

Samples were re-suspended in 4% formic acid, 2% acetonitrile solution, and separated by a reversed-phase gradient over a nanoflow C18 column (Dr. Maisch). Each sample was directly injected via a Easy-nLC 1200 (Thermo Fisher Scientific) into a Q-Exactive Plus mass spectrometer (Thermo Fisher Scientific) and analyzed with a 75 min acquisition, with all MS1 and MS2 spectra collected in the orbitrap; data were acquired using the Thermo software Xcalibur (4.2.47) and Tune (2.11 QF1 Build 3006). For all acquisitions, QCloud was used to control instrument longitudinal performance during the project^53^. All proteomic data was searched against the human proteome (uniprot reviewed sequences downloaded February 28th, 2020), EGFP sequence, and the SARS-CoV-2 protein sequences using the default settings for MaxQuant (version 1.6.12.0)^54,55^. Detected peptides and proteins were filtered to 1% false discovery rate in MaxQuant.

Identified proteins were then subjected to protein-protein interaction scoring with both SAINTexpress (version 3.6.3)^56^ and MiST (https://github.com/kroganlab/mist) ^57,58^. Interactions passing the master threshold (MiST score ≥ 0.6, a SAINTexpress BFDR ≤ 0.05 and an average spectral count ≥ 2) for at least one of the baits (mutants or wild-types) were kept for further analysis. An “Interaction Score” was defined as the average between the MiST score and the Saint Score, as previously described^10^. In addition, interactions were removed if their detection was found to be discrepant for wild-type Nsp2 between this study and a prior study^2^ (difference in interaction scores between studies < 0.4), further increasing the confidence of our final set of interactions. Remaining interactions were separated into two groups: those interacting with Nsp2 wild-type or not (average Interaction Score > 0.5 or < 0.5 were separated into Fig 5A-C and Sup Fig 5A, respectively). For those interacting with wild-type, a quantitative statistical analysis was performed (this quantitation was not possible for those not interacting with wild-type Nsp2 due to the lack of detection). Specifically, prey intensities in each affinity purification were normalized to the corresponding bait abundance using MSstats^59^ (globalStandards norm). Log_2_ fold changes and BH-adjusted p-values were calculated by comparing each mutant to the wild-type from this study.

#### Scheduled parallel reaction monitoring (PRM) analysis of Nsp2 interactors

Peptides for targeted MS were selected after importing the msms.txt file derived from the previously described MaxQuant search into Skyline (v20.2.0.343)^60^. Proteotypic peptides passing an Andromeda score of 0.95 were selected and manually inspected to choose precursors suitable for targeted proteomics. In total 4 peptides per protein were selected for targeted analysis. For WASHC4 and EIF4E2 all peptides identified by DDA were used (3 and 2 respectively). The samples from AP-MS were acquired in Partial Reaction Monitoring mode (PRM)^61^ on a Q-Exactive Orbitrap (Thermo Fisher) connected to a nanoLC easy 1200 (Thermo Fisher). Peptides for the scheduled analysis were separated in 75 minutes using the same chromatographic gradient and source parameter to the DDA samples. Precursor ion scans were recorded in the Orbitrap at 70’000 resolution (at 400 m/z) for 100 ms or until the ion population reached an AGC value of 1e^6^. Peptides in the inclusion list were fragmented using HCD with a normalized collisional energy of 27, an isolation window of 2 Da and a scheduled retention time window of 7 minutes. Fragments were acquired in the Orbitrap at 17’500 resolution (at 400 m/z) for 100 ms or until reaching an AGC of 2e^5^. Loop count was set to 20. For data analysis, the PRM data was searched with MaxQuant using a FASTA file containing only the target proteins and default settings. The msms.txt was imported into Skyline using the ‘import peptide search’ option and setting the search type to targeted. To import the files, the following transition settings were used: The MS1 filter was disabled, ion types were set to y and b and MS/MS settings were set to Orbitrap as mass analyzer, type as targeted and resolution of 17500 (at 400 m/z). Peptides with poor coeluting fragments (dotp lower than 0.9) were removed. WASHC4 peptides did not pass this quality control criterion and thus WASHC4 was not considered for further analysis. After import, peak boundaries were manually corrected and noisy transitions were removed. The resulting data was exported at the transition level and transitions missing in more than half of the samples were removed to ensure robust quantitation. The resulting transitions were summed up per peptide and then the experiment was mean centered using the average peptide level for the bait protein (using SARS-Cov-2 Nsp2 quantity for SARS-CoV-2 mutants). Normalized peptide-level abundances were averaged to reach protein level and log_2_ transformed. The fold change and BH-adjusted p-values for condition were calculated against the wild-type Nsp2.

## Supporting information

Supplemental figures and table 1 with cryoEM/model statistics

Table with the thresholded and un-thresholded mass spectrometry results.

## Acknowledgments

The authors acknowledge their partners and families for support in childcare and other matters during this time. We thank D. Mullins for helpful discussions and V. Ramani for providing critical feedback on the manuscript. The structural biology portion of this work was performed by the QCRG (<u>Q</u>uantitative Biosciences Institute <u>C</u>oronavirus Research <u>G</u>roup) Structural Biology Consortium. Listed below are the contributing members of the consortium listed by teams in order of team relevance to the published work. Within each team the team leads are italicized (responsible for organization of each team, and for the experimental design utilized within each team), then the rest of team members are listed alphabetically. **Bacterial expression team**: *Amy Diallo, Meghna Gupta, Erron W. Titus*, Jen Chen, Loan Doan, Sebastian Flores, Mingliang Jin, Huong T. Kratochvil, Victor L. Lam, Yang Li, Megan Lo, Gregory E. Merz, Joana Paulino, Aye C. Thwin, Zanlin Yu, Fengbo Zhou, Yang Zhang. **Protein purification team**: *Michelle Moritz, Tristan W. Owens, Sergei Pourmal*, Caleigh M. Azumaya, Cynthia M. Chio, Bryan Faust, Meghna Gupta, Kate Kim, Joana Paulino, Komal Ishwar Pawar, Jessica K. Peters, Kaitlin Schaefer, Ursula Schulze-Gahmen, Tsz Kin Martin Tsui. **CryoEM grid freezing/collection team:** *Caleigh M. Azumaya, Axel F. Brilot, Gregory E. Merz, Cristina Puchades, Alexandrea N. Rizo, Ming Sun*, Julian R. Braxton, Meghna Gupta, Fei Li, Kyle E. Lopez, Arthur Melo, Gregory E. Merz, Frank R. Moss III, Joana Paulino, Thomas H. Pospiech, Jr., Sergei Pourmal, Amber M. Smith, Paul V. Thomas, Feng Wang, Zanlin Yu. **CryoEM data processing team:** *Axel F. Brilot, Miles Sasha Dickinson, Gregory E. Merz, Henry C. Nguyen, Alexandrea N. Rizo*, Daniel Asarnow, Julian R. Braxton, Melody G. Campbell, Cynthia M. Chio, Un Seng Chio, Devan Diwanji, Bryan Faust, Soumya Govinda Remesh, Meghna Gupta, Nick Hoppe, Mingliang Jin, Fei Li, Junrui Li, Yanxin Liu, Adamo Mancino, Melissa Mendez, Joana Paulino, Thomas H. Pospiech, Jr., Sergei Pourmal, Smriti Sangwan, Raphael Trenker, Donovan Trinidad, Eric Tse, Kaihua Zhang, Fengbo Zhou. **Mammalian cell expression team:** *Christian Billesboelle, Melody G. Campbell, Devan Diwanji, Evelyn Hernandez, Carlos Nowotny, Amber M. Smith, Jianhua Zhao*, Caleigh M. Azumaya, Alisa Bowen, Nick Hoppe, Yen-Li Li, Edmond Linossi, Jocelyne Lopez, Phuong Nguyen, Michael D. Paul, Cristina Puchades, Mali Safari, Smriti Sangwan, Kaitlin Schaefer, Raphael Trenker, Tsz Kin Martin Tsui, Natalie Whitis. **Crystallography team:** *Nadia Herrera, Huong T. Kratochvil, Ursula Schulze-Gahmen, Iris D. Young*, Justin Biel, Ishan Deshpande, Xi Liu. **Infrastructure team:** David Bulkley, Arceli Joves, Almarie Joves, Liam McKay, Mariano C. Tabios, Eric Tse. **Leadership team:** *Oren S Rosenberg, Kliment A Verba*, David A Agard, Yifan Cheng, James S Fraser, Adam Frost, Natalia Jura, Tanja Kortemme, Nevan J Krogan, Aashish Manglik, Daniel R. Southworth, Robert M Stroud. The QCRG Structural Biology Consortium has received support from: Quantitative Biosciences Institute, Defense Advance Research Projects Agency HR0011-19-2-0020 (to D.A.A. and K.A.V.; B. Shoichet PI), FastGrants COVID19 grant (K.A.Verba PI), Laboratory For Genomics Research (O.S. Rosenberg PI) and Laboratory For Genomics Research (R.M. Stroud PI)

## Author contribution statement

The following authors designed and generated protein constructs and also performed protein expressions: M.G., A.D., G.E.M., D.C.D., E.H., V.L.L., Y.L., C.A.N., A.M.S., Z.Y., M.G.C., J.C., L.D., Y.L., E.L., M.L., J.L., M.D.P., M.S., A.C.T., E.W.T., R.T., Y.Z., J.Z.. The following authors performed protein purifications: C.M.A., M.M., S.P., T.W.O., J.K.P., U.S.G., K.K., K.I.P., K.S., T.K.M.T., F.Z.. The following authors performed structural analysis (data collection by crystallography or cryoEM and data processing and model building): M.G., C.M.A., S.P., G.E.M., A.F.B., N.H., H.T.K., F.L., H.C.N., A.N.R., I.D.Y., D.E.A., U.S.C., M.S.D., M.J., J.L., Y.L., K.E.L., A.M., F.R.M., T.H.P.Jr, C.P., S.G.R., M.S., E.T.. Graphene oxide grids were provided by F.W. and Z.Y.. M.C.T. provided crucial infrastructural and organizational support. The following authors did affinity purification mass spectrometry studies: G.M.J., M.B., A.F., K.C., A.P., Y.Z.. The following authors supervised or managed research: D.A.A., Y.C., J.S.F., N.J., T.K., A.M., D.R.S., R.M.S., D.L.S., N.J.K., A.F., O.S.R., K.A.V.. The following authors designed and conceptualized the study: M.G., C.M.A., M.M., S.P., A.D., G.E.M., M.B., L.Z.A., D.A.A., Y.C., J.S.F., N.J., T.K., A.M., D.R.S., R.M.S., N.J.K., A.F., O.S.R., K.A.V.. The following drafted the initial manuscript: M.G., M.M., S.P., G.E.M., G.M.J., M.B., A.F., F.L., L.Z.A., J.S.F., D.L.S., N.J.K., A.F., O.S.R., K.A.V.. All authors edited the manuscript.

