## Supplemental figures and table 1 with cryoEM/model statistics for "CryoEM and AI reveal a structure of SARS-CoV-2 Nsp2, a multifunctional protein involved in key host processes"

**Supplementary figure 1. Multiple sequence alignment of Nsp2.** Conserved cysteines are marked with red arrows (see positions 230-260 in the alignment).

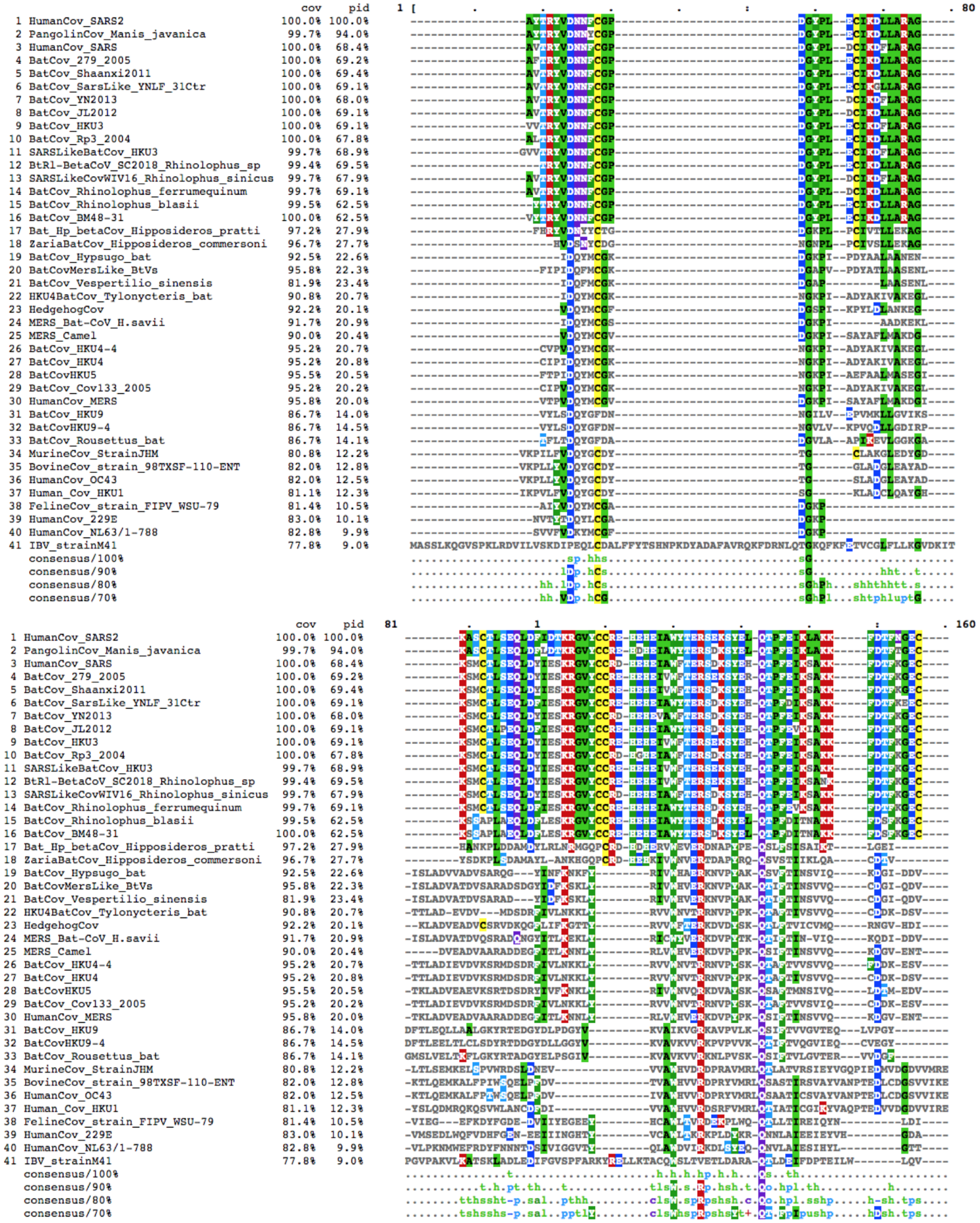

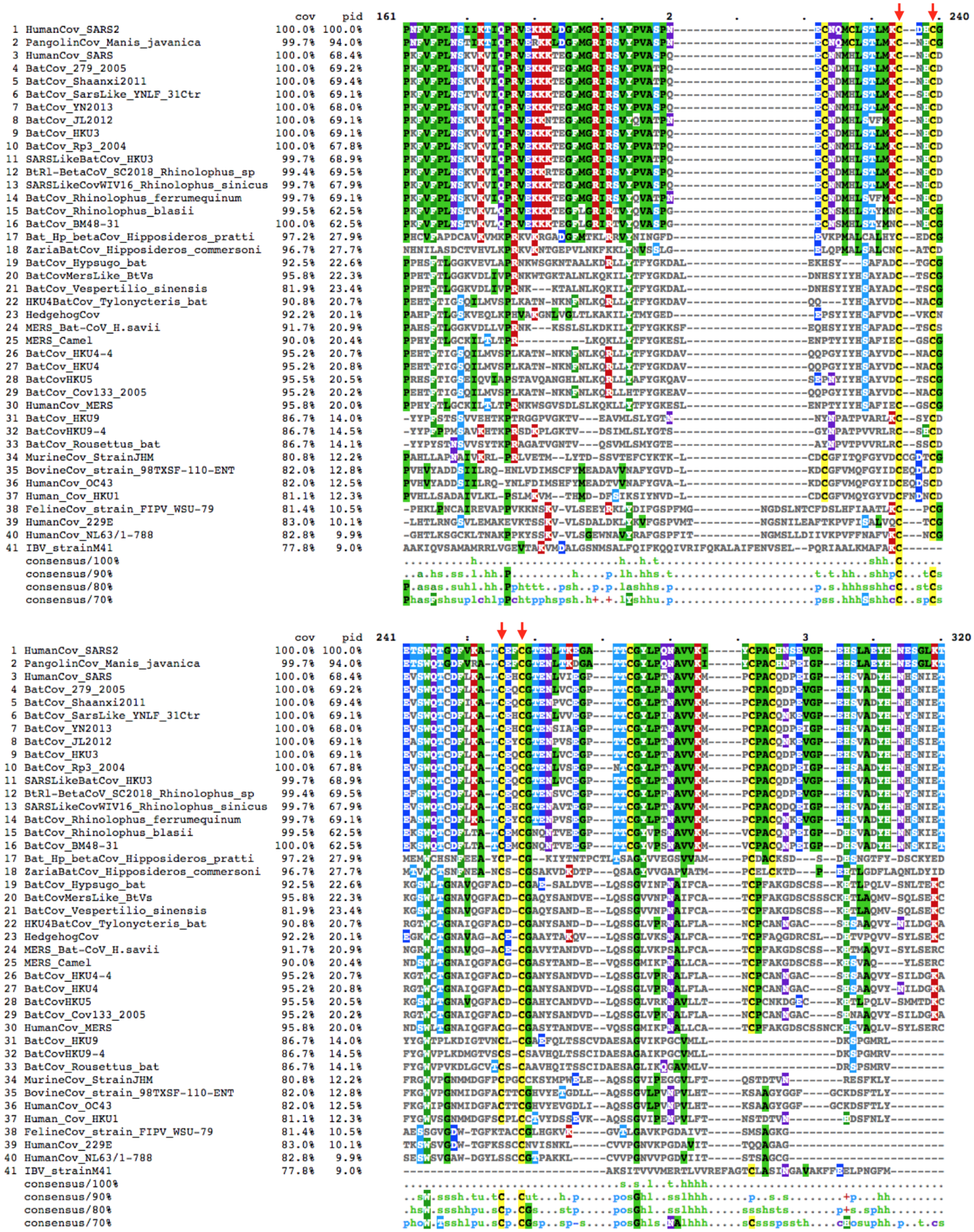

cov
pid
161
240
241
320
1 HumanCov_SARS2
2 PangolinCov_Manis_javanica
3 HumanCov_SARS
4 BatCov_279_2005
5 BatCov_Shaanxi2011
6 BatCov_SarsLike_YNLF_31Ctr
7 BatCov_YN2013
8 BatCov_JL2012
9 BatCov_HKU3
10 BatCov_Rp3_2004
11 SARSLikeBatCov_HKU3
12 BtRl-BetaCoV_SC2018_Rhinolophus_sp
13 SARSLikeCovWIV16_Rhinolophus_sinicus
14 BatCov_Rhinolophus_ferrumequinum
15 BatCov_Rhinolophus_blasii
16 BatCov_BM48-31
17 Bat_Hp_betaCov_Hipposideros_pratti
18 ZariaBatCov_Hipposideros_commersoni
19 BatCov_Hypsugo_bat
20 BatCovMersLike_BtVs
21 BatCov_Vespertilio_sinensis
22 HKU4BatCov_Tylonycteris_bat
23 HedgehogCov
24 MERS_Bat-CoV_H.savii
25 MERS_Camel
26 BatCov_HKU4-4
27 BatCov_HKU4
28 BatCovHKU5
29 BatCov_Cov133_2005
30 HumanCov_MERS
31 BatCov_HKU9
32 BatCovHKU9-4
33 BatCov_Rousettus_bat
34 MurineCov_StrainJHM
35 BovineCov_strain_98TXSF-110-ENT
36 HumanCov_OC43
37 Human_Cov_HKU1
38 FelineCov_strain_FIPV_WSU-79
39 HumanCov_229E
40 HumanCov_NL63/1-788
41 IBV_strainM41
consensus/100%
consensus/90%
consensus/80%
consensus/70%

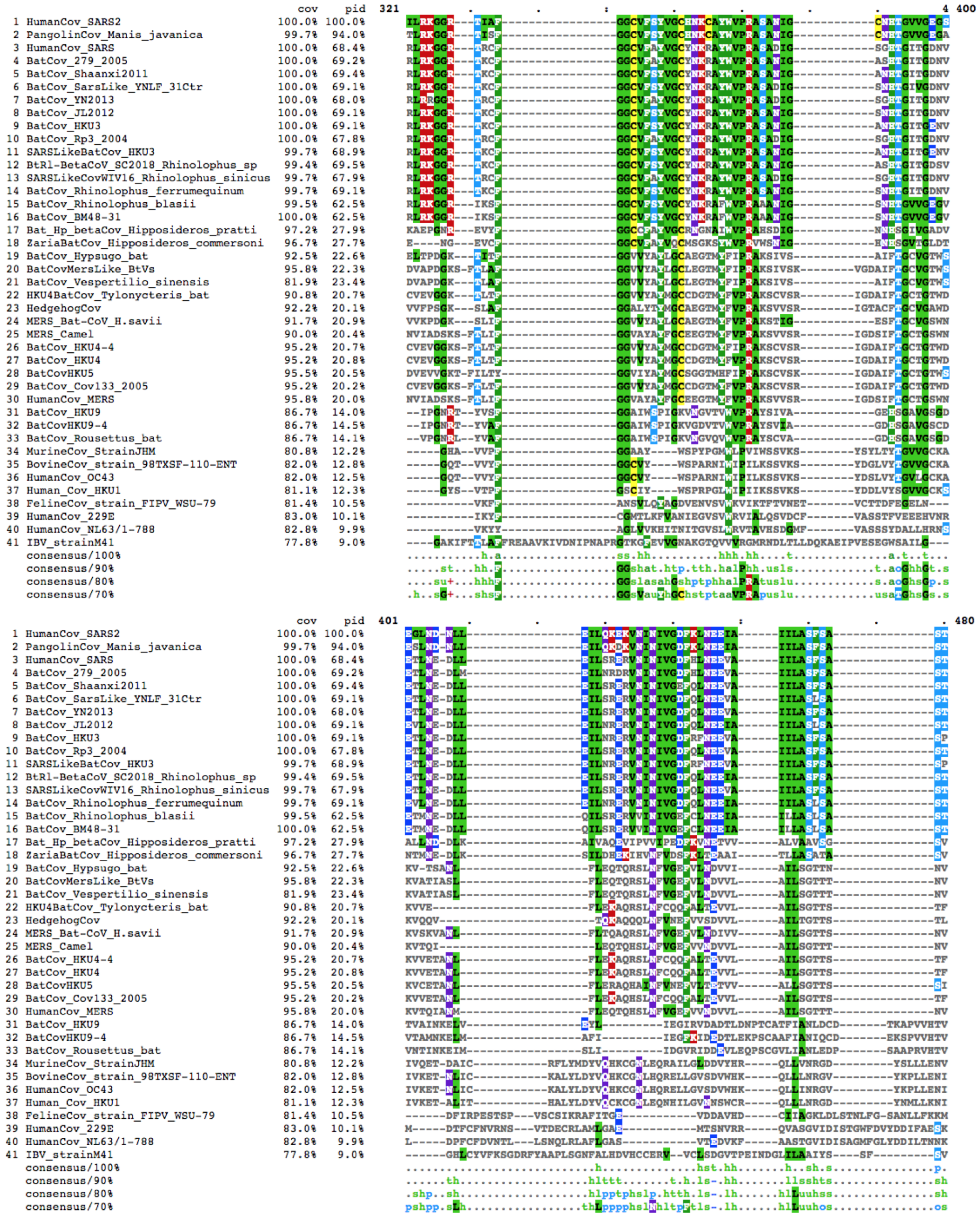
cov pid 321 400
1 HumanCov_SARS2 100.0% 100.0%
2 PangolinCov_Manis_javanica 99.7% 94.0%
3 HumanCov_SARS 100.0% 68.4%
4 BatCov_279_2005 100.0% 69.2%
5 BatCov_Shaanxi2011 100.0% 69.4%
6 BatCov_SarsLike_YNLF_31Ctr 100.0% 69.1%
7 BatCov_YN2013 100.0% 68.0%
8 BatCov_JL2012 100.0% 69.1%
9 BatCov_HKU3 100.0% 69.1%
10 BatCov_Rp3_2004 100.0% 67.8%
11 SARSLikeBatCov_HKU3 99.7% 68.9%
12 BtRl-BetaCoV_SC2018_Rhinolophus_sp 99.4% 69.5%
13 SARSLikeCovWIV16_Rhinolophus_sinicus 99.7% 67.9%
14 BatCov_Rhinolophus_ferrumequinum 99.7% 69.1%
15 BatCov_Rhinolophus_blasii 99.5% 62.5%
16 BatCov_BM48-31 100.0% 62.5%
17 Bat_Hp_betaCov_Hipposideros_pratti 97.2% 27.9%
18 ZariaBatCov_Hipposideros_commersoni 96.7% 27.7%
19 BatCov_Hypsugo_bat 92.5% 22.6%
20 BatCovMersLike_BtVs 95.8% 22.3%
21 BatCov_Vespertilio_sinensis 81.9% 23.4%
22 HKU4BatCov_Tylonycteris_bat 90.8% 20.7%
23 HedgehogCov 92.2% 20.1%
24 MERS_Bat-CoV_H.savii 91.7% 20.9%
25 MERS_Camel 90.0% 20.4%
26 BatCov_HKU4-4 95.2% 20.7%
27 BatCov_HKU4 95.2% 20.8%
28 BatCovHKU5 95.5% 20.5%
29 BatCov_Cov133_2005 95.2% 20.2%
30 HumanCov_MERS 95.8% 20.0%
31 BatCov_HKU9 86.7% 14.0%
32 BatCovHKU9-4 86.7% 14.5%
33 BatCov_Rousettus_bat 86.7% 14.1%
34 MurineCov_StrainJHM 80.8% 12.2%
35 BovineCov_strain_98TXSF-110-ENT 82.0% 12.8%
36 HumanCov_OC43 82.0% 12.5%
37 Human_Cov_HKU1 81.1% 12.3%
38 FelineCov_strain_FIPV_WSU-79 81.4% 10.5%
39 HumanCov_229E 83.0% 10.1%
40 HumanCov_NL63/1-788 82.8% 9.9%
41 IBV_strainM41 77.8% 9.0%
consensus/100%
consensus/90%
consensus/80%
consensus/70%
cov pid 401 480
1 HumanCov_SARS2 100.0% 100.0%
2 PangolinCov_Manis_javanica 99.7% 94.0%
3 HumanCov_SARS 100.0% 68.4%
4 BatCov_279_2005 100.0% 69.2%
5 BatCov_Shaanxi2011 100.0% 69.4%
6 BatCov_SarsLike_YNLF_31Ctr 100.0% 69.1%
7 BatCov_YN2013 100.0% 68.0%
8 BatCov_JL2012 100.0% 69.1%
9 BatCov_HKU3 100.0% 69.1%
10 BatCov_Rp3_2004 100.0% 67.8%
11 SARSLikeBatCov_HKU3 99.7% 68.9%
12 BtRl-BetaCoV_SC2018_Rhinolophus_sp 99.4% 69.5%
13 SARSLikeCovWIV16_Rhinolophus_sinicus 99.7% 67.9%
14 BatCov_Rhinolophus_ferrumequinum 99.7% 69.1%
15 BatCov_Rhinolophus_blasii 99.5% 62.5%
16 BatCov_BM48-31 100.0% 62.5%
17 Bat_Hp_betaCov_Hipposideros_pratti 97.2% 27.9%
18 ZariaBatCov_Hipposideros_commersoni 96.7% 27.7%
19 BatCov_Hypsugo_bat 92.5% 22.6%
20 BatCovMersLike_BtVs 95.8% 22.3%
21 BatCov_Vespertilio_sinensis 81.9% 23.4%
22 HKU4BatCov_Tylonycteris_bat 90.8% 20.7%
23 HedgehogCov 92.2% 20.1%
24 MERS_Bat-CoV_H.savii 91.7% 20.9%
25 MERS_Camel 90.0% 20.4%
26 BatCov_HKU4-4 95.2% 20.7%
27 BatCov_HKU4 95.2% 20.8%
28 BatCovHKU5 95.5% 20.5%
29 BatCov_Cov133_2005 95.2% 20.2%
30 HumanCov_MERS 95.8% 20.0%
31 BatCov_HKU9 86.7% 14.0%
32 BatCovHKU9-4 86.7% 14.5%
33 BatCov_Rousettus_bat 86.7% 14.1%
34 MurineCov_StrainJHM 80.8% 12.2%
35 BovineCov_strain_98TXSF-110-ENT 82.0% 12.8%
36 HumanCov_OC43 82.0% 12.5%
37 Human_Cov_HKU1 81.1% 12.3%
38 FelineCov_strain_FIPV_WSU-79 81.4% 10.5%
39 HumanCov_229E 83.0% 10.1%
40 HumanCov_NL63/1-788 82.8% 9.9%
41 IBV_strainM41 77.8% 9.0%
consensus/100%
consensus/90%
consensus/80%
consensus/70%

| | | cov | pid | 481 . 5 . . . . : . 560 |
|---|---|---|---|---|
| 1 | HumanCov_SARS2 | 100.0% | 100.0% | SAFVETVKGLDYKAFKQIVESCGNFKVTKGKAKKGAWNIGEQ----KSILSPLYAFA--SEAARVVRSIFSRTLETAQNS |
| 2 | PangolinCov_Manis_javanica | 99.7% | 94.0% | SAFVETVKGLDYKTFKQIVESCGNFKVTKGKAKKGAWNIGEQ----KSILSPLYAFA--SEAARVIRAIFSRTLETAQHS |
| 3 | HumanCov_SARS | 100.0% | 68.4% | SAFIDTIKSLDYKSFKTIVESCGNYKVTKGKPVKGAWNIGQQ----RSVLTPLCGFP--SQAAGVIRSIFARTLDAANHS |
| 4 | BatCov_279_2005 | 100.0% | 69.2% | CAFVDTVKGLDYKTFKDIVESCGNFKVTRGRAKKGAWNIGQE----KSILTPLYGFP--SQAAGVIRSIFTRALDTANHS |
| 5 | BatCov_Shaanxi2011 | 100.0% | 69.4% | SAFIDTVKSLDYKSFKAIVESCGNYKVTKGKPVKGAWNIGQQ----RSILTPLCGFP--SQAAGVIRSIFSRTLDAANHS |
| 6 | BatCov_SarsLike_YNLF_31Ctr | 100.0% | 69.1% | SAFIDTIKSLDYKSFKSIVESCGNYKVTKGKPIKGAWNIGQQ----RSVLTPLCGFP--SQAAGVIRSIFARTLDAANHS |
| 7 | BatCov_YN2013 | 100.0% | 68.0% | SAFIDTIKSLDYKSFKSIVESCGNYKVTKGKPIKGAWNIGQQ----RSVLTPLCGFP--SQAAGVIRSIFARTLDAANHS |
| 8 | BatCov_JL2012 | 100.0% | 69.1% | SAFVDTVKSLDYKSFKAIVESCGNYKVTKGKPVKSAWNIGQQ----KSILTPLCGFP--SQAAGVIRSIFSRTLDAANNS |
| 9 | BatCov_HKU3 | 100.0% | 69.1% | SAFIETVKGLDYKSFKVIVESCGNYKVTNGKPVTGAWNIGQQ----RSILTPLCGFP--SQAAGVIRSIFSRTLDAANHS |
| 10 | BatCov_Rp3_2004 | 100.0% | 67.8% | SAFIDTIKSLDYKSFKTIVESCGNYKVTKGKPIKGAWNIGQH----RSVLTPLCGFP--SQAAGVIRSIFSRTLDAANHS |
| 11 | SARSLikeBatCov_HKU3 | 99.7% | 68.9% | SAFIETVKGLDYKSFKVIVESCGNYKVTNGKPVTGAWNIGQQ----RSILTPLCGFP--SQAAGVIRSIFSRTLDAANHS |
| 12 | BtRl-BetaCoV_SC2018_Rhinolophus_sp | 99.4% | 69.5% | SAFIDTVKSLDYKSFKAIVESCGNYKVTKGKPVKGAWNIGQQ----KSILTPLCGFP--SQAAGVIRSIFSRTLDAANHS |
| 13 | SARSLikeCovWIV16_Rhinolophus_sinicus | 99.7% | 67.9% | SAFIDTIRSLDYKSFKAIVESCGNYKVTKGKPVKGAWNIGQQ----RSVLTPLCGFP--SQAAGVIRSIFARTLDAANHS |
| 14 | BatCov_Rhinolophus_ferrumequinum | 99.7% | 69.1% | SAFVDTVKSLDYKSFKAIVESCGNYKVTKGKPVKGAWNIGQQ----KSILTPLCGFP--SQAAGVTRSIFSRTLDAANNS |
| 15 | BatCov_Rhinolophus_blasii | 99.5% | 62.5% | SAFVETVKNLDFKTFKKIIESCGNYKVTKGKFKPGVWNIGTS----KSLLTPLHCFS--SQAAGVVRSIFSRTLATANHS |
| 16 | BatCov_BM48-31 | 100.0% | 62.5% | SAFVETVKNLDFKTFKKIIESCGNYKVTKGKFKPGVWNIGTS----KSLLTPLHCFS--SQAAGVVRSIFSRTLATANHS |
| 17 | Bat_Hp_betaCov_Hipposideros_pratti | 97.2% | 27.9% | DQFIKNCKGLTAETFKCLLEASGKFRVTVDHVIDGCICVATD----SAVLEPALAFC--TAANIMYRTLVATSVVISAHT |
| 18 | ZariaBatCov_Hipposideros_commersoni | 96.7% | 27.7% | ESLIQNVKDLTFDKYKILLESTGRFQVTMGKYVQGALNFSKE----DTFCDPILAAP--GHALSLCRAIFNKTVDLSGTT |
| 19 | BatCov_Hypsugo_bat | 92.5% | 22.6% | DKLRSLVSNLDADKLRNYLSDFG-IAVTVGPYLDGAINVGKQ----ATITAPFL-------SFKKAAAVPYKS--SLKDT |
| 20 | BatCovMersLike_BtVs | 95.8% | 22.3% | DKLRNLVKHLDVDKLREYLADYD-IAVTVGPYLDGAINVGKQGLQFAAITAPFVVLTGLGESFKKVAAIPYKACKSIKET |
| 21 | BatCov_Vespertilio_sinensis | 81.9% | 23.4% | DKLRNLVKHLDVDKLREYLAGYD-IAVTVGPYLDGAINVGKQ----AAITAPFVVLT--GESFKKVAAIPYKACKSIKET |
| 22 | HKU4BatCov_Tylonycteris_bat | 90.8% | 20.7% | EELRDLCHNASYEKVRDHLVN---FVVTIGDYIRDAINIGA----NATINAPFIAFT--GESFKKVSAIPWKICSNLKSA |
| 23 | HedgehogCov | 92.2% | 20.1% | DELRSLLQGITFEKMRDYC--HG-IKVTMGDYVDSAINVGRS----AAINAPFVVLSALGESFKKIAAVPFKVGSSFLKT |
| 24 | MERS_Bat-CoV_H.savii | 91.7% | 20.9% | EKLRSLIQDISFERLRDYLAAYD-IEVTKGSFVEGAINVGKN----AAITAPFLVLL--AESFKKVAAVPYKVCCTLRDT |
| 25 | MERS_Camel | 90.0% | 20.4% | DKIRQLLKGVTLDKLRDYLADYD-VVVTAGPFMDNAINVG----QYAAITAPYVVLT--GESFKKVATIPYKVCYSVKDT |
| 26 | BatCov_HKU4-4 | 95.2% | 20.7% | EELHDLCHNASYEKVRDHLVNHG-FVVTIGDYIRDAINIGANGVCNATINAPFIAFTGLGESFKKVSAIPWKICSNLKSA |
| 27 | BatCov_HKU4 | 95.2% | 20.8% | EELRDLCHNASYEKVRDHLVNHG-FVVTIGDYIRDAINIGANGVCNATINAPFIAFTGLGESFKKVSAIPWKICSNLKSA |
| 28 | BatCovHKU5 | 95.5% | 20.5% | EELRDLCRNATFEKVRDYLTPRG-WIVTMGSYIEGVINVGAAGVCNAALNAPFIVLSGLGESFKKVAATPWKLCSSLRET |
| 29 | BatCov_Cov133_2005 | 95.2% | 20.2% | EELRDLCHNASYEKVRDHLVNHG-FVVTIGDYIRDAINIGANGVCNATINAPFIAFTGLGESFKKVAAIPWKICSNLKSA |
| 30 | HumanCov_MERS | 95.8% | 20.0% | DKIRQLLKGVTLDKLRDYLADYD-VAVTAGPFMDNAINVGGTGLQYAAITAPYVVLTGLGESFKKVATIPYKVCNSVKDT |
| 31 | BatCov_HKU9 | 86.7% | 14.0% | ESLQGLC-------------------------LANKIMLGDKPLPTDEFHPFIVGLAYHVQRACWYGALASRTFEAFRDF |
| 32 | BatCovHKU9-4 | 86.7% | 14.5% | DSITKMC-------------------------LDNKVVLGEHPLPDDEFHPAIVGMNYYTQRTCWYAALSSKTFAAMRDY |
| 33 | BatCov_Rousettus_bat | 86.7% | 14.1% | DSLRQLC-------------------------VDNNVMLGDTPLPKDEFHPGIVGLSYHFYRACWYGVLTAKSFGAFKEL |
| 34 | MurineCov_StrainJHM | 80.8% | 12.2% | DLFVKRRAEFACK-FA--TCGDGLVPLLLDGLVPRSYYLIKS-----------------GQAFTSLMV------------ |
| 35 | BovineCov_strain_98TXSF-110-ENT | 82.0% | 12.8% | DYFNMRRAKFSLETFT--VCADGFMPFLLDDLVPRAYYLAVS-----------------GQAFCDYAD------------ |
| 36 | HumanCov_OC43 | 82.0% | 12.5% | DYFNMRRAKFSLETFT--VCADGFMPFLLDDLVPRAYYLAVS-----------------GQAFCDYAD------------ |
| 37 | Human_Cov_HKU1 | 81.1% | 12.3% | DLFVKRRADFACK-FA--VCGDGFVPFLLDGLIPRSYYLIQS-----------------GIFFTSLMS------------ |
| 38 | FelineCov_strain_FIPV_WSU-79 | 81.4% | 10.5% | PWFVQKCGAIFADAWKVVEELLCSLKLTYKQIYDVVASLCTS---AFTIMDYKPVFV--VS-SNSVKDLVDKCVKILVKA |
| 39 | HumanCov_229E | 83.0% | 10.1% | PWFVRKAEDIFGPCWSALASALKQLKVTTGELVRFVKSICNS---AVAVVGGTIQIL--ASVPEKFLNAFDVFVTAIQTV |
| 40 | HumanCov_NL63/1-788 | 82.8% | 9.9% | PWFVRKASGLFDAIWDAFVAAIKLVPTTTGVLVRFVKSIAST---VLTVSNGVIIMC--ADVPDAFQSVYRTFTQAICAA |
| 41 | IBV_strainM41 | 77.8% | 9.0% | AELVAAIKRGEPFKFLG-------HKFVYAKDAAVSFTLAKA----ATI----------ADVLKLFQSARVKV------- |
| | consensus/100% | | | ..h...h.........................h...h.h..........................h.s............ |
| | consensus/90% | | | p.hhphhtth....ht........h.hh.s..h.tsh.lutp.....th..........sp.ht.htsh..h........ |
| | consensus/80% | | | sthhphhpthshcphp.hl.s.s.h.lThs..hpsshslutp....tsl.sshhshs..upuhthhtsl..+hhtshtps |
| | consensus/70% | | | sthhphhcshshcph+thltshs.hhVThGp.lcsuhNlGpp....tslhsPhhshs..upuhthltul..+shpshpps |

| | | cov | pid | 561 . . . 6 . . . . 640 |
|---|---|---|---|---|
| 1 | HumanCov_SARS2 | 100.0% | 100.0% | VRVLQKAAI----------------------------------------------------TILDGISQYSLRLIDAMM- |
| 2 | PangolinCov_Manis_javanica | 99.7% | 94.0% | VRVLQQAAI----------------------------------------------------TILDGISQYSLRLIDAMM- |
| 3 | HumanCov_SARS | 100.0% | 68.4% | IPDLQRAAV----------------------------------------------------TILDGISEQSLRLVDAMV- |
| 4 | BatCov_279_2005 | 100.0% | 69.2% | IPDLQRAAI----------------------------------------------------TILDGISEQSLRLIDAMV- |
| 5 | BatCov_Shaanxi2011 | 100.0% | 69.4% | IPDLQRAAV----------------------------------------------------TILDGISEQSLRLVDAMV- |
| 6 | BatCov_SarsLike_YNLF_31Ctr | 100.0% | 69.1% | IPDLQRAAV----------------------------------------------------TILGGISEQSLRLVDAMV- |
| 7 | BatCov_YN2013 | 100.0% | 68.0% | IPDLQRAAV----------------------------------------------------TILDGISEQSLRLVDAMV- |
| 8 | BatCov_JL2012 | 100.0% | 69.1% | IPDLQRAAV----------------------------------------------------TILGGISEQSLRLVDAMV- |
| 9 | BatCov_HKU3 | 100.0% | 69.1% | ILDLQRAAV----------------------------------------------------TTLDGISEQSLRLVDAMV- |
| 10 | BatCov_Rp3_2004 | 100.0% | 67.8% | IPDLQRAAV----------------------------------------------------TILDSISEQSLRLVDAMV- |
| 11 | SARSLikeBatCov_HKU3 | 99.7% | 68.9% | ILDLQRAAV----------------------------------------------------TTLDGISEQSLRLVDAMV- |
| 12 | BtRl-BetaCoV_SC2018_Rhinolophus_sp | 99.4% | 69.5% | IPDLQRAAV----------------------------------------------------TILDGISEQSLRLVDAMV- |
| 13 | SARSLikeCovWIV16_Rhinolophus_sinicus | 99.7% | 67.9% | IPDLQRAAV----------------------------------------------------TILDGISEQSLRLVDAMV- |
| 14 | BatCov_Rhinolophus_ferrumequinum | 99.7% | 69.1% | IPDLQRAAV----------------------------------------------------TILGDISEQSLRLVDAMV- |
| 15 | BatCov_Rhinolophus_blasii | 99.5% | 62.5% | IVDLHRAAM----------------------------------------------------IIFSDISDQANRVLDAMV- |
| 16 | BatCov_BM48-31 | 100.0% | 62.5% | IVDLHRAAM----------------------------------------------------IIFSDISDQANRVLDAMV- |
| 17 | Bat_Hp_betaCov_Hipposideros_pratti | 97.2% | 27.9% | LDALAQRVR----------------------------------------------------DACKHVHPQAMRLLNTLN- |
| 18 | ZariaBatCov_Hipposideros_commersoni | 96.7% | 27.7% | LANLKETMS----------------------------------------------------AVYKDLHPRALKLVCTLC- |
| 19 | BatCov_Hypsugo_bat | 92.5% | 22.6% | LDYYANSILYRVFP----------------------------------------------YEISSDVSSFSELLLDC--- |
| 20 | BatCovMersLike_BtVs | 95.8% | 22.3% | LLYYAESILYRVFP----------------------------------------------YEISSDVSTFSELLLDC--- |
| 21 | BatCov_Vespertilio_sinensis | 81.9% | 23.4% | LLYYAESIL----------------------------------------------------EISSDVSTFSELLLDC--- |
| 22 | HKU4BatCov_Tylonycteris_bat | 90.8% | 20.7% | LDYYSSNIMFRVFP----------------------------------------------YDIPCDVSNFVELLLDC--- |
| 23 | HedgehogCov | 92.2% | 20.1% | WEYLSDCIVYRVLP----------------------------------------------YELE-DV-DFVQLLFNC--- |
| 24 | MERS_Bat-CoV_H.savii | 91.7% | 20.9% | LEYYAQSIL----------------------------------------------------DMSSDVSSFSELLLEC--- |
| 25 | MERS_Camel | 90.0% | 20.4% | L-----SVLYRVFP----------------------------------------------YDMD-GVSSFSELLFDC--- |
| 26 | BatCov_HKU4-4 | 95.2% | 20.7% | LDYYSSNIMFRVFP----------------------------------------------YDIPCDVSNFVELLLDC--- |
| 27 | BatCov_HKU4 | 95.2% | 20.8% | LDYYSSNIMFRVFP----------------------------------------------YDIPCDVSNFVELLLDC--- |
| 28 | BatCovHKU5 | 95.5% | 20.5% | LDHYADSITYRVFP----------------------------------------------YDIPCDVTDYTALLLDC--- |
| 29 | BatCov_Cov133_2005 | 95.2% | 20.2% | LDYYCSNIMFRVFP----------------------------------------------YDIPCDVNDFVELLLDC--- |
| 30 | HumanCov_MERS | 95.8% | 20.0% | LTYYAHSVLYRVFP----------------------------------------------YDMDSGVSSFSELLFDC--- |
| 31 | BatCov_HKU9 | 86.7% | 14.0% | VRTEEERFA----------------------------------------------------QFFGKVCAPINGCVYLAY- |
| 32 | BatCovHKU9-4 | 86.7% | 14.5% | VRREEERLR----------------------------------------------------VFCGKAFTPVSACLQMAY- |
| 33 | BatCov_Rousettus_bat | 86.7% | 14.1% | LQSEEVRLS----------------------------------------------------HFCANIRRCLDRALNWAR- |
| 34 | MurineCov_StrainJHM | 80.8% | 12.2% | -------------------------------------------------------------NFSREVVDMCMDMALLFM- |
| 35 | BovineCov_strain_98TXSF-110-ENT | 82.0% | 12.8% | -------------------------------------------------------------KICHAVVSKSKELLDVSL- |
| 36 | HumanCov_OC43 | 82.0% | 12.5% | -------------------------------------------------------------KLCHAVVSKSKELLDVSL- |
| 37 | Human_Cov_HKU1 | 81.1% | 12.3% | -------------------------------------------------------------QFSQEVSDMCLKMCILFM- |
| 38 | FelineCov_strain_FIPV_WSU-79 | 81.4% | 10.5% | FDVFTQTITIAGVEAKCFVLGSKYLLFNNALVKLVSVKILGKRQKGLDSAFFATNLIGATVNVTPQRTESAYISLNKVD- |
| 39 | HumanCov_229E | 83.0% | 10.1% | FDCAVETCTIAGKA---FDKVFDYVLLDNALVKLVTTKLKGVRERGLNKVKYATVVVGSTEEVKSSRVERSTAVLTIANN |
| 40 | HumanCov_NL63/1-788 | 82.8% | 9.9% | FDFSLDVFKIGDVK---FKRLGDYVLTENALVRLTTEVVRGVRDARIKKAMFTKVVVGPTTEVKFSVIELATVNLRLVDC |
| 41 | IBV_strainM41 | 77.8% | 9.0% | ------------------------------------------------------------EDVWSSLTEKSFEFWRLAY- |
| | consensus/100% | | | ..............................................................h..th.......h.h... |
| | consensus/90% | | | .............................................................ph.ttl.p.s.hhhph... |
| | consensus/80% | | | h...tpthh....................................................ph.tslsp.s.hllsh... |
| | consensus/70% | | | l..htpshh....................................................pl.sslop.u.hLl-s... |

```
                                          cov    pid  641        :         .         .         .         .         7         .         . 720
 1 HumanCov_SARS2                       100.0% 100.0%      ------------------------------FTSDLATNNLVVMAYIT-------------GGVVQLTSQWLTNI------
 2 PangolinCov_Manis_javanica            99.7%  94.0%      ------------------------------FTSDLVTDNLVVMAYIT-------------GGVVQMTSQWLTNI------
 3 HumanCov_SARS                        100.0%  68.4%      ------------------------------YTSDLLTNSVIIMAYVT-------------GGLVQQTSQWLSNL------
 4 BatCov_279_2005                      100.0%  69.2%      ------------------------------YTSDLLTNSVIVMAYVT-------------GGLVQQITQWLSNM------
 5 BatCov_Shaanxi2011                   100.0%  69.4%      ------------------------------YTSDLITNSVIVMAYVT-------------GGLVQQTTQWLSNM------
 6 BatCov_SarsLike_YNLF_31Ctr           100.0%  69.1%      ------------------------------YTSDLLTNSVIVMAYVT-------------GGLVQQTSQWLSNL------
 7 BatCov_YN2013                        100.0%  68.0%      ------------------------------YTSDLLTNSVIIMAYVT-------------GGLVQQTSQWLSNL------
 8 BatCov_JL2012                        100.0%  69.1%      ------------------------------CTSDLITNSVVIMAYVT-------------GGLVQQTVQWLSNV------
 9 BatCov_HKU3                          100.0%  69.1%      ------------------------------YTSDLLTNSVVVMAYVT-------------GGLVQQTMQWLSNM------
10 BatCov_Rp3_2004                      100.0%  67.8%      ------------------------------YTSNLLTNSVIIMAYVT-------------GGLVQQTSQWLSNL------
11 SARSLikeBatCov_HKU3                   99.7%  68.9%      ------------------------------YTSDLLTNSVVVMAYVT-------------GGLVQQTMQWLSNM------
12 BtRl-BetaCoV_SC2018_Rhinolophus_sp    99.4%  69.5%      ------------------------------YTSDLLTNSVIVMAYVT-------------GGLVQQTTQWLSNM------
13 SARSLikeCovWIV16_Rhinolophus_sinicus  99.7%  67.9%      ------------------------------YTSDLITNSVIIMAYVT-------------GGLVQQISQWLSNL------
14 BatCov_Rhinolophus_ferrumequinum      99.7%  69.1%      ------------------------------CTSDLITNSVVIMAYVT-------------GGLLQQTVQWLSNV------
15 BatCov_Rhinolophus_blasii             99.5%  62.5%      ------------------------------NTSDLVTESVVVMAYLT-------------GGLVQQVSTWLSQL------
16 BatCov_BM48-31                       100.0%  62.5%      ------------------------------NTSDLVTESVVVMAYLT-------------GGLVQQVSTWLSQL------
17 Bat_Hp_betaCov_Hipposideros_pratti    97.2%  27.9%      ------------------------------EYVDYSTNSTLVLCHDV-------------INLPIVLGDILDRL------
18 ZariaBatCov_Hipposideros_commersoni   96.7%  27.7%      ------------------------------DNVTLMRNSVLLLMQDA-------------QNLPVVIGDMLDCL------
19 BatCov_Hypsugo_bat                    92.5%  22.6%      --------------------------------IGFTAASAYFVVRL----------------LHDKTDTLLSTI------
20 BatCovMersLike_BtVs                   95.8%  22.3%      --------------------------------VGFTAASCYFVVRL----------------LDEKAEALLSTV------
21 BatCov_Vespertilio_sinensis           81.9%  23.4%      --------------------------------VGFTAASCYFVVRL----------------LDEKAEALLSTV------
22 HKU4BatCov_Tylonycteris_bat           90.8%  20.7%      --------------------------------GKLTVATSYFVLRY----------------LDEKFDTVLGTV------
23 HedgehogCov                           92.2%  20.1%      --------------------------------VEISAASMYFASVV----------------IREKVNTMFNAL------
24 MERS_Bat-CoV_H.savii                  91.7%  20.9%      --------------------------------VGFTTASVYFVARL----------------LDEKVESLLSTI------
25 MERS_Camel                            90.0%  20.4%      --------------------------------VDLSVASTYFLVRL----------------LQ-KTGDFMSTI------
26 BatCov_HKU4-4                         95.2%  20.7%      --------------------------------GKLTVAISYFVLRY----------------LDEKFDTVLGTV------
27 BatCov_HKU4                           95.2%  20.8%      --------------------------------GKLTVATSYFVLRY----------------LDEKFDTVLGTV------
28 BatCovHKU5                            95.5%  20.5%      --------------------------------AVLTGASAYFVARY----------------VDEKVEQLTNLV------
29 BatCov_Cov133_2005                    95.2%  20.2%      --------------------------------GKLTVATSYFVLRY----------------LDEKFDTVLGTV------
30 HumanCov_MERS                         95.8%  20.0%      --------------------------------VDLSVASTYFLVRL----------------LQDKTGDFMSTI------
31 BatCov_HKU9                           86.7%  14.0%      ------------------------------TTGRVTLFSAYQVLNT----------------AIAKSKDAFGGV------
32 BatCovHKU9-4                          86.7%  14.5%      ------------------------------QSGRVTLVTAYQVLDT----------------ALTKTKDAFGGA------
33 BatCov_Rousettus_bat                  86.7%  14.1%      ------------------------------TTGNVSLLSSSLVLKA----------------LAENLLDSVKNT------
34 MurineCov_StrainJHM                   80.8%  12.2%      --------------------------------HDVKVATKYVKKVT------------------GKVAVRFKAL------
35 BovineCov_strain_98TXSF-110-ENT       82.0%  12.8%      --------------------------------DSLSAAIHYLNSKI------------------VDLAQHFSDF------
36 HumanCov_OC43                         82.0%  12.5%      --------------------------------DSLGAAIHYLNSKI------------------VDLAQHFSDF------
37 Human_Cov_HKU1                        81.1%  12.3%      --------------------------------DRVSVATFYIEHYV------------------NRLVTQFKLL------
38 FelineCov_strain_FIPV_WSU-79          81.4%  10.5%      -DVVTPGGGHIVIIGDMAFYKSEEYYFMMASPDSVLVNNVFKAARVPSYNIVYDVNDDTKSKMVVKIGTSFD-F------
39 HumanCov_229E                         83.0%  10.1%      YSKLFD-EGYTVVIGDVAYFVSDGYFRLMASPNSVLTTAVYKP------LFAFNVN---------VMGTRPEKF-PTTVT
40 HumanCov_NL63/1-788                   82.8%   9.9%      APVVCP-KGKIVVIAGQAFFYSGGFYRFMVDPTTVLNDPVFTG------DLFYTIK---------FSGFKLDGFNHQFVT
41 IBV_strainM41                         77.8%   9.0%      --------GKV-------------------RNLEEFVKTCFCKAQMA------------IVILATVLGEGIWHL------
   consensus/100%                                          .........................................................................h......
   consensus/90%                                           ................................sth.sts.hhh..h....................h.phhtth......
   consensus/80%                                           ................................splhssoshhhhhh................h.tphtphhssh......
   consensus/70%                                           ................................sslhssoshhhshh................l.pphsphlssl......

                                          cov    pid  721        .         .         :         .         .         .         .         8 800
 1 HumanCov_SARS2                       100.0% 100.0%      FGTVYEKLKPVLDWLEEKF-------------------------------------------------------------
 2 PangolinCov_Manis_javanica            99.7%  94.0%      FGTVYEKLKPILEWLEDKF-------------------------------------------------------------
 3 HumanCov_SARS                        100.0%  68.4%      LGTTVEKLRPIFEWIEAKL-------------------------------------------------------------
 4 BatCov_279_2005                      100.0%  69.2%      LGTTVDKLKPVFTWVEAKL-------------------------------------------------------------
 5 BatCov_Shaanxi2011                   100.0%  69.4%      LGTTVEKLKPVFTWIETKL-------------------------------------------------------------
 6 BatCov_SarsLike_YNLF_31Ctr           100.0%  69.1%      LGTTVEKLRPIFAWIEAKL-------------------------------------------------------------
 7 BatCov_YN2013                        100.0%  68.0%      LGTTVEKLRPIFSWIEAKL-------------------------------------------------------------
 8 BatCov_JL2012                        100.0%  69.1%      LGTTVDKLKPVFTWLETKL-------------------------------------------------------------
 9 BatCov_HKU3                          100.0%  69.1%      LGTAVDKLKPVFTWVEAKL-------------------------------------------------------------
10 BatCov_Rp3_2004                      100.0%  67.8%      LDTTVEKLRPIFAWIEAKL-------------------------------------------------------------
11 SARSLikeBatCov_HKU3                   99.7%  68.9%      LGTAVDKLKPVFTWVEAKL-------------------------------------------------------------
12 BtRl-BetaCoV_SC2018_Rhinolophus_sp    99.4%  69.5%      LGTTVDKLRPVFTWVETKL-------------------------------------------------------------
13 SARSLikeCovWIV16_Rhinolophus_sinicus  99.7%  67.9%      LGTTVEKLRPIFAWIEAKL-------------------------------------------------------------
14 BatCov_Rhinolophus_ferrumequinum      99.7%  69.1%      LGTTVDKLKPVFTWLETKL-------------------------------------------------------------
15 BatCov_Rhinolophus_blasii             99.5%  62.5%      LNTSVDKFSAVLRWLEQKL-------------------------------------------------------------
16 BatCov_BM48-31                       100.0%  62.5%      LNTSVDKFSAVLRWLEQKL-------------------------------------------------------------
17 Bat_Hp_betaCov_Hipposideros_pratti    97.2%  27.9%      FCGCKKIVESVTTWLYNFS-------------------------------------------------------------
18 ZariaBatCov_Hipposideros_commersoni   96.7%  27.7%      FQGCADFVNKVLSKLMEFA-------------------------------------------------------------
19 BatCov_Hypsugo_bat                    92.5%  22.6%      FSSCQAAVKSFLNACFETT-------------------------------------------------------------
20 BatCovMersLike_BtVs                   95.8%  22.3%      FSSCQAAVSKFLNGCFEAT-------------------------------------------------------------
21 BatCov_Vespertilio_sinensis           81.9%  23.4%      FSSCQAAVSKFLNGCFEAT-------------------------------------------------------------
22 HKU4BatCov_Tylonycteris_bat           90.8%  20.7%      SS----ALSSFLNACVAAS-------------------------------------------------------------
23 HedgehogCov                           92.2%  20.1%      PLSVQTAAKNFIDVCLRAT-------------------------------------------------------------
24 MERS_Bat-CoV_H.savii                  91.7%  20.9%      FSN-------FITTCFEAT-------------------------------------------------------------
25 MERS_Camel                            90.0%  20.4%      IISCQTAVSKFLDTCFEAT-------------------------------------------------------------
26 BatCov_HKU4-4                         95.2%  20.7%      SSACQTALSSFLNACVAAS-------------------------------------------------------------
27 BatCov_HKU4                           95.2%  20.8%      SSACQTALSSFLNACVAAS-------------------------------------------------------------
28 BatCovHKU5                            95.5%  20.5%      FSSCQSAVAAFVQACMSTY-------------------------------------------------------------
29 BatCov_Cov133_2005                    95.2%  20.2%      SNACQTALSSFLNACVAAS-------------------------------------------------------------
30 HumanCov_MERS                         95.8%  20.0%      ITSCQTAVSKLLDTCFEAT-------------------------------------------------------------
31 BatCov_HKU9                           86.7%  14.0%      AAIVVDMLKPILEWVLKKM-------------------------------------------------------------
32 BatCovHKU9-4                          86.7%  14.5%      AVVVSDLLKPTLDWVLTKM-------------------------------------------------------------
33 BatCov_Rousettus_bat                  86.7%  14.1%      PFLVGDLVKPVLDWIVEKM-------------------------------------------------------------
34 MurineCov_StrainJHM                   80.8%  12.2%      GIAVVRKITEWFDLAVDTA-------------------------------------------------------------
35 BovineCov_strain_98TXSF-110-ENT       82.0%  12.8%      GTSFVSKIVHFFKTFTTST-------------------------------------------------------------
36 HumanCov_OC43                         82.0%  12.5%      GTSFVSKIVHFFKTFTTST-------------------------------------------------------------
37 Human_Cov_HKU1                        81.1%  12.3%      GTTLVNKMVNWFNTMLDAS-------------------------------------------------------------
38 FelineCov_strain_FIPV_WSU-79          81.4%  10.5%      DGDLDAAIAKVNDLLIEFRQEKLCFRALKDGENILVEAYLKKYKMPVCLKNHVGLWD-IIRQDSGKKGFLDTFNHLNELE
39 HumanCov_229E                         83.0%  10.1%      CENLESAVLFVNDKITEFQ---LDYSIDVIDNEIIVKPNI-SLCVPLYVRDYVDKWDDFCRQYSNESWFEDDYRAFISVL
40 HumanCov_NL63/1-788                   82.8%   9.9%      ASSATDAIIAVELLLLDFKTAVFVYTCVVDGCSVIVRRDA-TFATHVCFKDCYNVWEQFCIDNCGEPWFLTDYNAILQSN
41 IBV_strainM41                         77.8%   9.0%      VSQVIYKVGGLFTKVVDFCE---------------------KY------------WKGFCAQ------------------
   consensus/100%                                          ..........h..hh.t...............................................................
   consensus/90%                                           ..sh.thh..hhphh.th..............................................................
   consensus/80%                                           hsss.ptlt.hhphh.tth.............................................................
   consensus/70%                                           hsss.ptlpshhshh.pth.............................................................
```

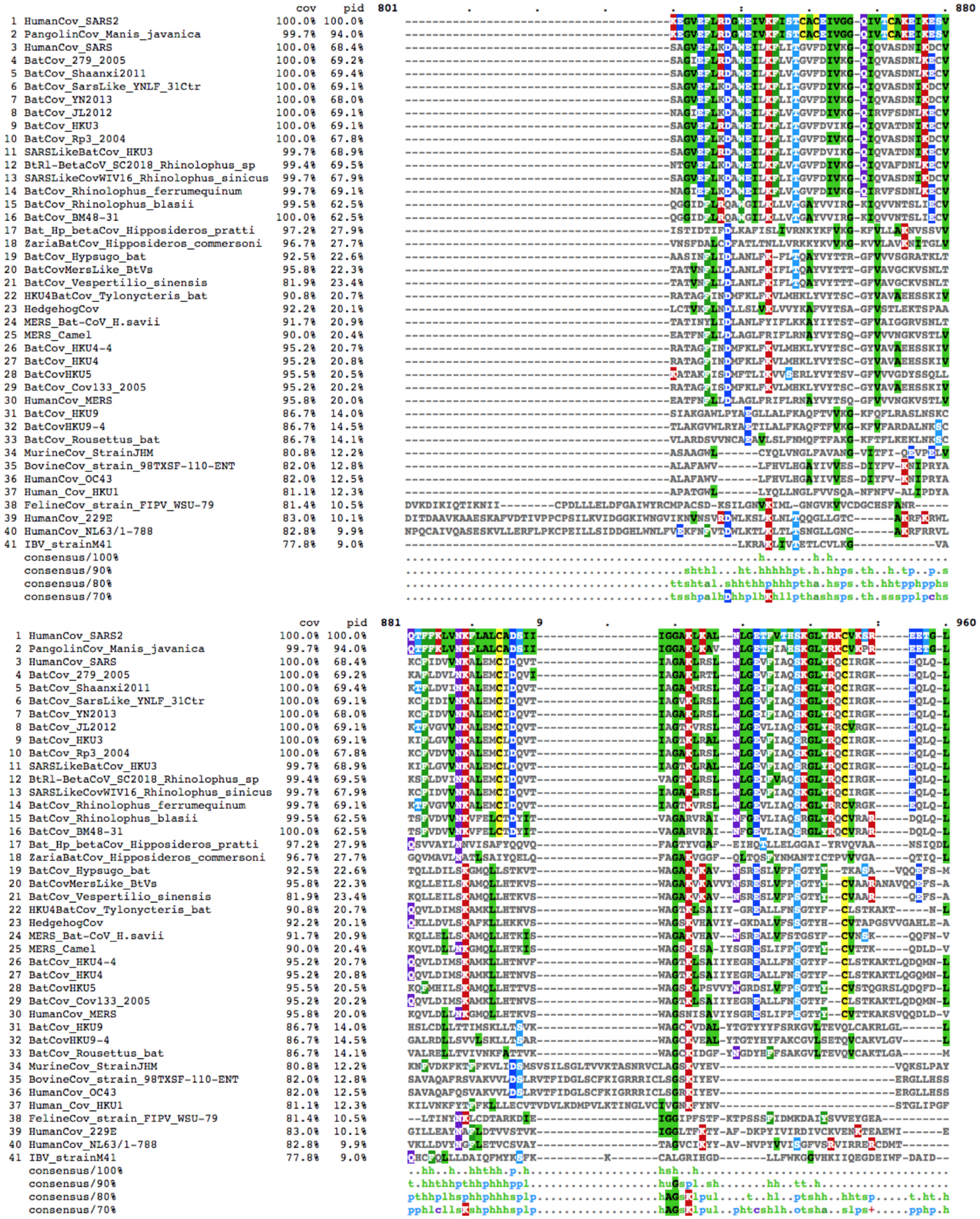
cov pid 801 880
1 HumanCov_SARS2 100.0% 100.0%
2 PangolinCov_Manis_javanica 99.7% 94.0%
3 HumanCov_SARS 100.0% 68.4%
4 BatCov_279_2005 100.0% 69.2%
5 BatCov_Shaanxi2011 100.0% 69.4%
6 BatCov_SarsLike_YNLF_31Ctr 100.0% 69.1%
7 BatCov_YN2013 100.0% 68.0%
8 BatCov_JL2012 100.0% 69.1%
9 BatCov_HKU3 100.0% 69.1%
10 BatCov_Rp3_2004 100.0% 67.8%
11 SARSLikeBatCov_HKU3 99.7% 68.9%
12 BtRl-BetaCoV_SC2018_Rhinolophus_sp 99.4% 69.5%
13 SARSLikeCovWIV16_Rhinolophus_sinicus 99.7% 67.9%
14 BatCov_Rhinolophus_ferrumequinum 99.7% 69.1%
15 BatCov_Rhinolophus_blasii 99.5% 62.5%
16 BatCov_BM48-31 100.0% 62.5%
17 Bat_Hp_betaCov_Hipposideros_pratti 97.2% 27.9%
18 ZariaBatCov_Hipposideros_commersoni 96.7% 27.7%
19 BatCov_Hypsugo_bat 92.5% 22.6%
20 BatCovMersLike_BtVs 95.8% 22.3%
21 BatCov_Vespertilio_sinensis 81.9% 23.4%
22 HKU4BatCov_Tylonycteris_bat 90.8% 20.7%
23 HedgehogCov 92.2% 20.1%
24 MERS_Bat-CoV_H.savii 91.7% 20.9%
25 MERS_Camel 90.0% 20.4%
26 BatCov_HKU4-4 95.2% 20.7%
27 BatCov_HKU4 95.2% 20.8%
28 BatCovHKU5 95.5% 20.5%
29 BatCov_Cov133_2005 95.2% 20.2%
30 HumanCov_MERS 95.8% 20.0%
31 BatCov_HKU9 86.7% 14.0%
32 BatCovHKU9-4 86.7% 14.5%
33 BatCov_Rousettus_bat 86.7% 14.1%
34 MurineCov_StrainJHM 80.8% 12.2%
35 BovineCov_strain_98TXSF-110-ENT 82.0% 12.8%
36 HumanCov_OC43 82.0% 12.5%
37 Human_Cov_HKU1 81.1% 12.3%
38 FelineCov_strain_FIPV_WSU-79 81.4% 10.5%
39 HumanCov_229E 83.0% 10.1%
40 HumanCov_NL63/1-788 82.8% 9.9%
41 IBV_strainM41 77.8% 9.0%
consensus/100%
consensus/90%
consensus/80%
consensus/70%
cov pid 881 960
1 HumanCov_SARS2 100.0% 100.0%
2 PangolinCov_Manis_javanica 99.7% 94.0%
3 HumanCov_SARS 100.0% 68.4%
4 BatCov_279_2005 100.0% 69.2%
5 BatCov_Shaanxi2011 100.0% 69.4%
6 BatCov_SarsLike_YNLF_31Ctr 100.0% 69.1%
7 BatCov_YN2013 100.0% 68.0%
8 BatCov_JL2012 100.0% 69.1%
9 BatCov_HKU3 100.0% 69.1%
10 BatCov_Rp3_2004 100.0% 67.8%
11 SARSLikeBatCov_HKU3 99.7% 68.9%
12 BtRl-BetaCoV_SC2018_Rhinolophus_sp 99.4% 69.5%
13 SARSLikeCovWIV16_Rhinolophus_sinicus 99.7% 67.9%
14 BatCov_Rhinolophus_ferrumequinum 99.7% 69.1%
15 BatCov_Rhinolophus_blasii 99.5% 62.5%
16 BatCov_BM48-31 100.0% 62.5%
17 Bat_Hp_betaCov_Hipposideros_pratti 97.2% 27.9%
18 ZariaBatCov_Hipposideros_commersoni 96.7% 27.7%
19 BatCov_Hypsugo_bat 92.5% 22.6%
20 BatCovMersLike_BtVs 95.8% 22.3%
21 BatCov_Vespertilio_sinensis 81.9% 23.4%
22 HKU4BatCov_Tylonycteris_bat 90.8% 20.7%
23 HedgehogCov 92.2% 20.1%
24 MERS_Bat-CoV_H.savii 91.7% 20.9%
25 MERS_Camel 90.0% 20.4%
26 BatCov_HKU4-4 95.2% 20.7%
27 BatCov_HKU4 95.2% 20.8%
28 BatCovHKU5 95.5% 20.5%
29 BatCov_Cov133_2005 95.2% 20.2%
30 HumanCov_MERS 95.8% 20.0%
31 BatCov_HKU9 86.7% 14.0%
32 BatCovHKU9-4 86.7% 14.5%
33 BatCov_Rousettus_bat 86.7% 14.1%
34 MurineCov_StrainJHM 80.8% 12.2%
35 BovineCov_strain_98TXSF-110-ENT 82.0% 12.8%
36 HumanCov_OC43 82.0% 12.5%
37 Human_Cov_HKU1 81.1% 12.3%
38 FelineCov_strain_FIPV_WSU-79 81.4% 10.5%
39 HumanCov_229E 83.0% 10.1%
40 HumanCov_NL63/1-788 82.8% 9.9%
41 IBV_strainM41 77.8% 9.0%
consensus/100%
consensus/90%
consensus/80%
consensus/70%

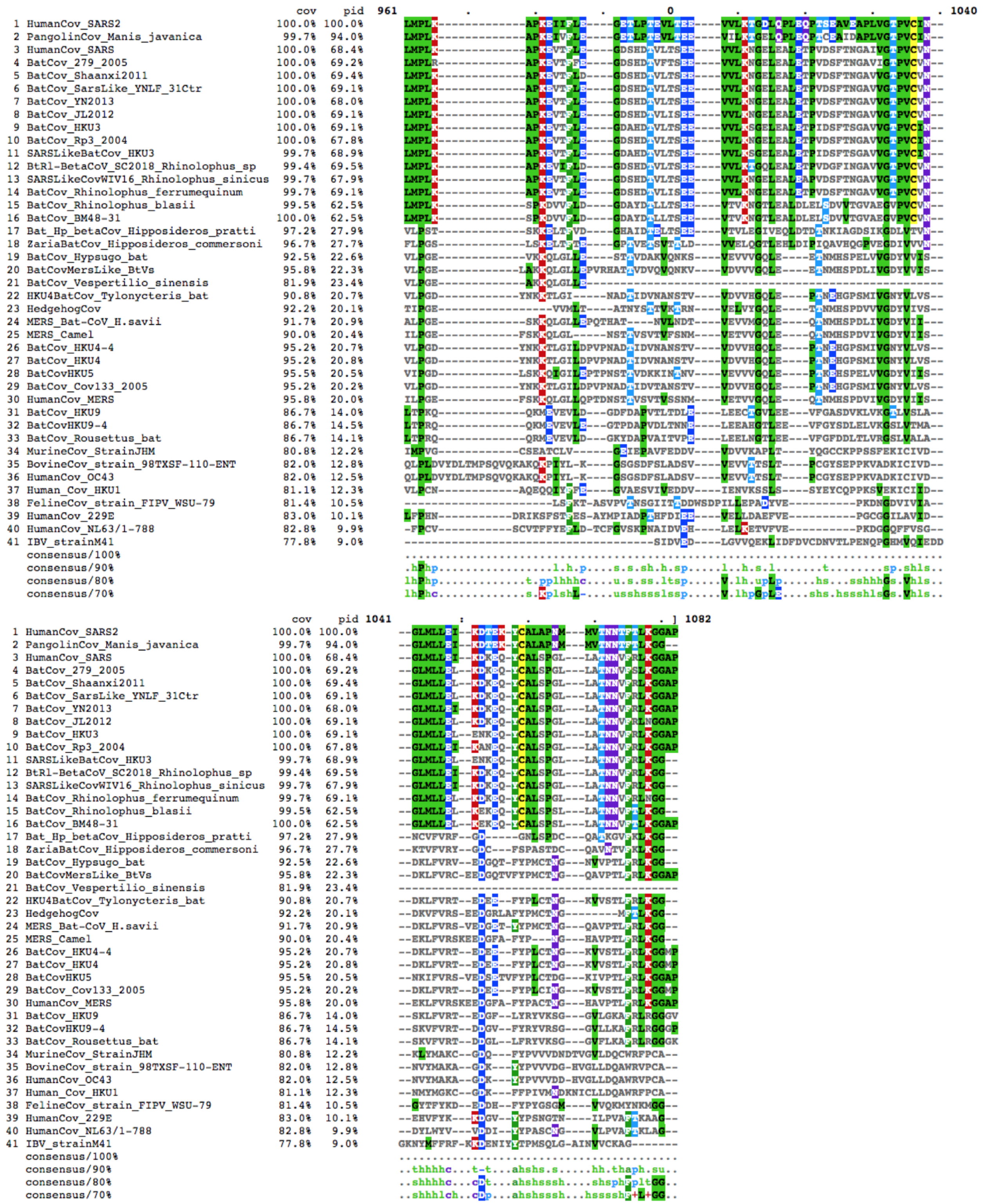
cov pid 961 1040
1 HumanCov_SARS2 100.0% 100.0%
2 PangolinCov_Manis_javanica 99.7% 94.0%
3 HumanCov_SARS 100.0% 68.4%
4 BatCov_279_2005 100.0% 69.2%
5 BatCov_Shaanxi2011 100.0% 69.4%
6 BatCov_SarsLike_YNLF_31Ctr 100.0% 69.1%
7 BatCov_YN2013 100.0% 68.0%
8 BatCov_JL2012 100.0% 69.1%
9 BatCov_HKU3 100.0% 69.1%
10 BatCov_Rp3_2004 100.0% 67.8%
11 SARSLikeBatCov_HKU3 99.7% 68.9%
12 BtRl-BetaCoV_SC2018_Rhinolophus_sp 99.4% 69.5%
13 SARSLikeCovWIV16_Rhinolophus_sinicus 99.7% 67.9%
14 BatCov_Rhinolophus_ferrumequinum 99.7% 69.1%
15 BatCov_Rhinolophus_blasii 99.5% 62.5%
16 BatCov_BM48-31 100.0% 62.5%
17 Bat_Hp_betaCov_Hipposideros_pratti 97.2% 27.9%
18 ZariaBatCov_Hipposideros_commersoni 96.7% 27.7%
19 BatCov_Hypsugo_bat 92.5% 22.6%
20 BatCovMersLike_BtVs 95.8% 22.3%
21 BatCov_Vespertilio_sinensis 81.9% 23.4%
22 HKU4BatCov_Tylonycteris_bat 90.8% 20.7%
23 HedgehogCov 92.2% 20.1%
24 MERS_Bat-CoV_H.savii 91.7% 20.9%
25 MERS_Camel 90.0% 20.4%
26 BatCov_HKU4-4 95.2% 20.7%
27 BatCov_HKU4 95.2% 20.8%
28 BatCovHKU5 95.5% 20.5%
29 BatCov_Cov133_2005 95.2% 20.2%
30 HumanCov_MERS 95.8% 20.0%
31 BatCov_HKU9 86.7% 14.0%
32 BatCovHKU9-4 86.7% 14.5%
33 BatCov_Rousettus_bat 86.7% 14.1%
34 MurineCov_StrainJHM 80.8% 12.2%
35 BovineCov_strain_98TXSF-110-ENT 82.0% 12.8%
36 HumanCov_OC43 82.0% 12.5%
37 Human_Cov_HKU1 81.1% 12.3%
38 FelineCov_strain_FIPV_WSU-79 81.4% 10.5%
39 HumanCov_229E 83.0% 10.1%
40 HumanCov_NL63/1-788 82.8% 9.9%
41 IBV_strainM41 77.8% 9.0%
consensus/100%
consensus/90%
consensus/80%
consensus/70%
cov pid 1041 1082

**Supplementary Figure 2.**

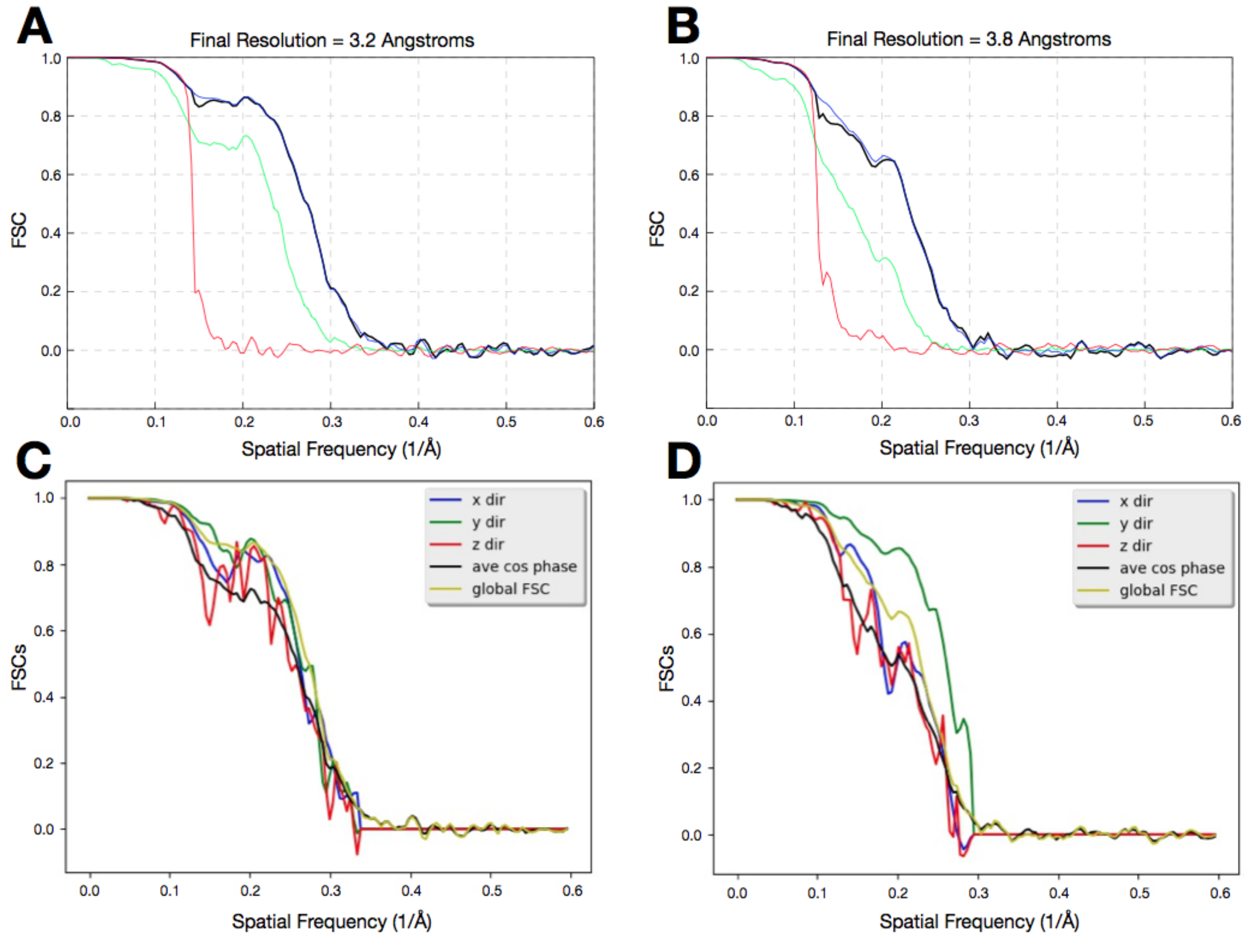


**Supplementary Figure 2. Global and directional FSC curves for the Nsp2 cryo-EM reconstructions.**

**(A)** Global FSC curve for the 3.2 Å Nsp2 reconstruction as output from Relion. In black is the final corrected masked FSC. In blue is an uncorrected masked FSC. In green is FSC of unmasked volumes and in red is FSC of the volumes which were phase randomized beyond 7.3 Å. **(B)** Global FSC curve for the 3.8 Å Nsp2 reconstruction as output from Relion. In black is the final corrected masked FSC. In blue is an uncorrected masked FSC. In green is unmasked FSC and in red is FSC of the reconstructions which were phase randomized beyond 8 Å. **(C)** Directional FSC for the 3.2 Å reconstruction of Nsp2 as output by the 3DFSC server, estimated sphericity is 0.86. **(D)** Directional FSC for the 3.8 Å reconstruction of Nsp2 as output by the 3DFSC server, estimated sphericity is 0.81.

**Supplementary Figure 3.**

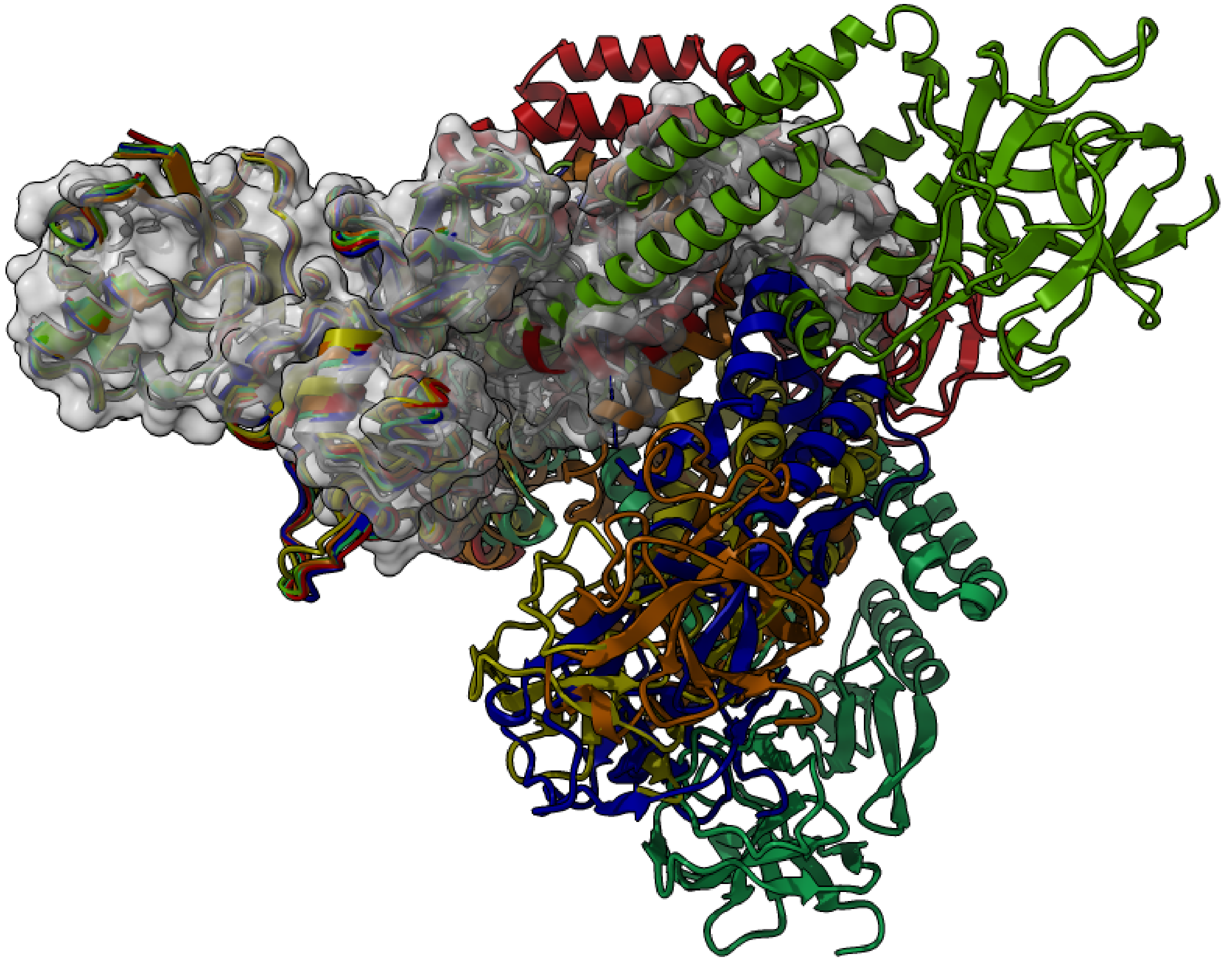

**Supplementary Figure 3. AlphaFold2 predictions for Nsp2 have low accuracy in predicting the overall shape of the protein.**

All publicly available predictions from AlphaFold2 team for Nsp2 (multicolored) were aligned to the experimental model of Nsp2 (gray surface). Although individual domains align well locally, globally there is a considerable deviation from the experimental model.

**Supplementary figure 4.**

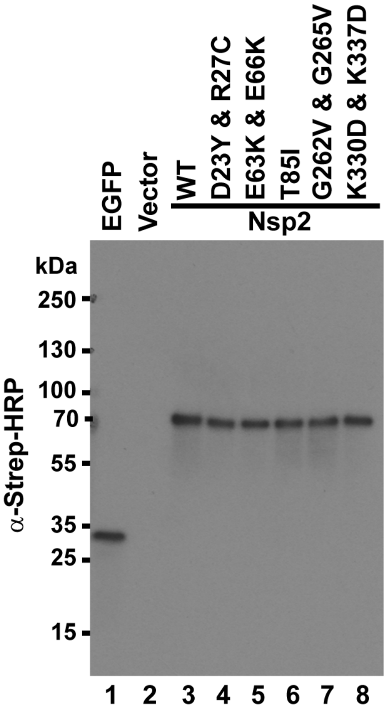


**Supplementary figure 4. Expression of wild type and mutant SARS-CoV-2 Nsp2 proteins in HEK293T cells.** Reserved lysates (50 ul) were incubated at 95°C for 5 minutes with an equal volume of 2x sample buffer (Morganville Scientific). After further diluting 1:10 in 2x sample buffer, 5 ul was resolved on a 4-20% Criterion TGX Precast Midi Protein Gel (BioRad) and transferred to a 0.2 mm PVDF membrane using the Trans-Blot Turbo Transfer System (2.5 A, 25 V, 7 minutes). Membranes were blocked with 3% BSA in 1x Phosphate Buffered Saline (PBS) supplemented with 0.2% Tween 20 (0.2% T-PBS) at 4°C and incubated with Strep-Tag II Antibody HRP Conjugate (Millipore) at room temperature for 1 hour (1:10,000 in 1% BSA, 0.2% T-PBS). Membrane was washed in 0.2% T-PBS after blocking and antibody incubation steps and developed with Pierce ECL Western Blotting Substrate (ThermoFisher Scientific).

### Supplementary figure 5.

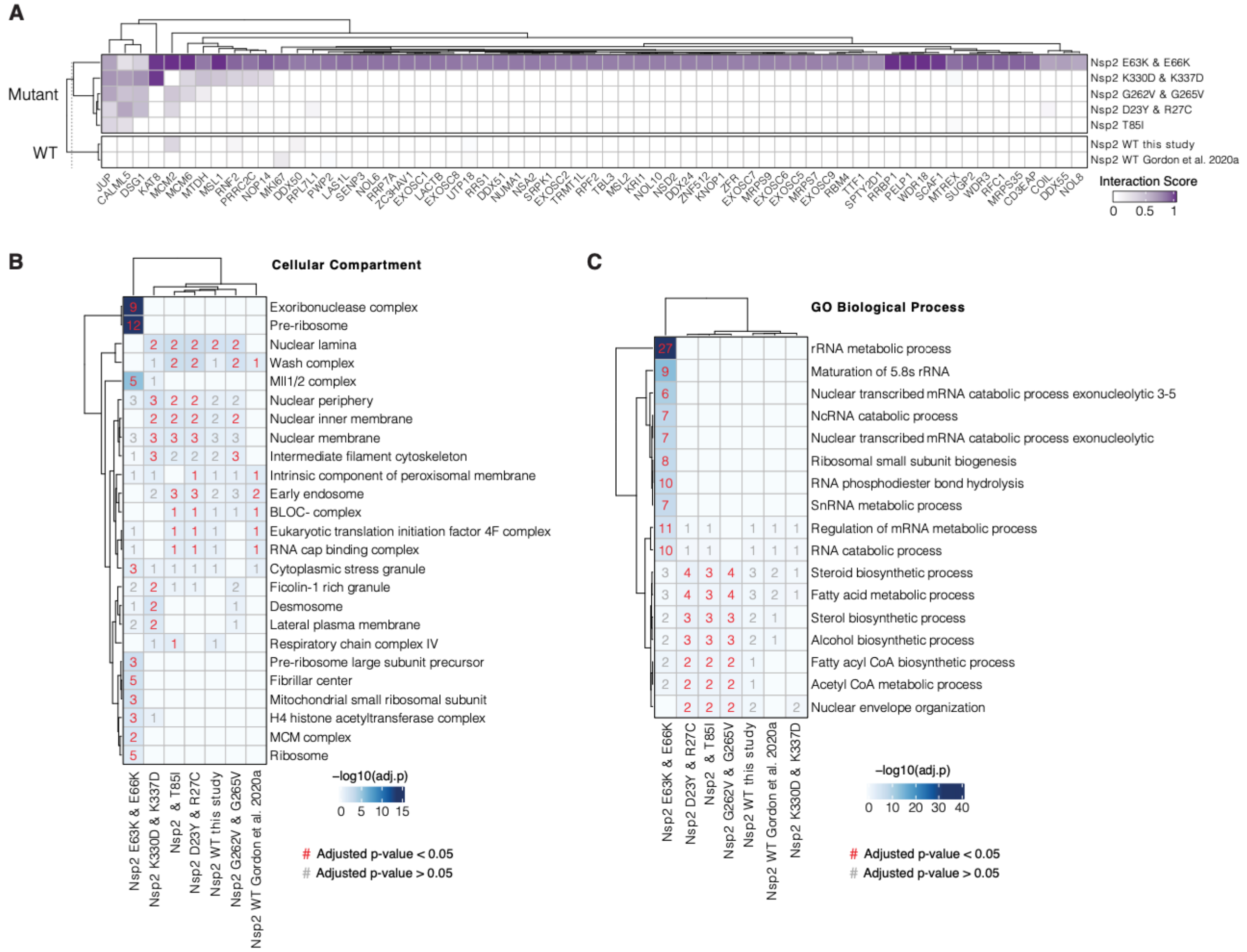


**Supplementary figure 5. Nsp2 E63K/E66K Mutation Gains Interactions with Exoribonuclease and Pre-ribosome Components.**

**(A)** Interaction scores (average between MiST and Saint Scores) for human proteins ("preys") deemed high-confidence interactions in at least one affinity purification ("bait") mass spectrometry assay and detected to not interact with neither the wild-type Nsp2 in this study or in Gordon et al (2020a). Interaction scores range from zero to one, one being the most high-confidence. **(B)** Gene Ontology Cellular Compartment enrichment analysis (MSigDB) for preys passing the master scoring criteria (see Methods) for each bait (i.e. affinity purification experiment). The top 10 most significant enrichments for each bait are displayed as well as corresponding enrichments for other baits, if applicable. Color denotes the -log$_{10}$(adjusted p-value). The number of preys enriched in each bait for each term are shown. Red numbers indicated adjusted p-values < 0.05 whereas grey numbers indicate adjusted p-values > 0.05. **(C)** Same as in B using Gene Ontology Biological Process terms (MSigDB).

**Supplementary Table 1. CryoEM collection, refinement and resulting model statistics.**

| | Nsp2 without Zn<br>(EMDB-xxxx)<br>(PDB xxxx) | Nsp2 with Zn<br>(EMDB-xxxx)<br>(PDB xxxx) |
|---|---|---|
| **Data collection and processing** | | |
| Magnification | 105,000x | 105,000x |
| Voltage (kV) | 300 | 300 |
| Electron exposure (e–/Å$^2$) | 66 | 67 |
| Dose rate (e-/physical pixel/sec) | 8 | 8 |
| Exposure per frame (sec) | 0.05 | 0.05 |
| Defocus range (μm) | -0.8 to -2.4 | -0.8 to -2.4 |
| Pixel size (Å) | 0.834 (physical) | 0.834 (physical) |
| Symmetry imposed | C1 | C1 |
| Initial particle images (no.) | 363145+577518 | 1515264 |
| Final particle images (no.) | 42579 | 81817 |
| Map resolution (Å)<br>FSC threshold | 3.76 | 3.15 |
| Map resolution range (Å) | 3.5-6.6 | 3.0-4.3 |
| **Refinement** | | |
| Initial model used (PDB code) | ab-initio | ab-initio |
| Model resolution (Å)<br>FSC threshold 0.143 (Unmasked)<br>FSC threshold 0.5 (Unmasked) | <br>3.6<br>3.9 | <br>3.12<br>3.46 |
| Model resolution range (Å) | 3.6-4.1 | 3.1-3.5 |
| Map sharpening *B* factor (Å$^2$) | -122 | -135 |
| Model composition<br>Non-hydrogen atoms<br>Protein residues<br>Ligands | <br>4922<br>635<br>Zn: 3 | <br>3706<br>473<br>Zn: 3 |
| *B* factors (Å$^2$)<br>Protein<br>Ligand | <br>194.4<br>79.8 | <br>52.83<br>50.97 |

| | | |
|---|---|---|
| R.M.S. deviations<br>Bond lengths (Å)<br>Bond angles (°) | <br>0.009<br>0.982 | <br>0.008<br>0.946 |
| Validation<br>MolProbity score<br>Clashscore<br>Poor rotamers (%) | <br>0.86<br>0.71<br>0.00 | <br>0.75<br>0.81<br>0.00 |
| Ramachandran plot<br>Favored (%)<br>Allowed (%)<br>Disallowed (%) | <br>97.31<br>2.53<br>0.16 | <br>98.07<br>1.93<br>0.00 |
